# Emergence of rare carbapenemases (FRI, GES-5, IMI, SFC, and SFH-1) in Enterobacterales isolated from surface waters in Japan

**DOI:** 10.1101/2021.10.04.462962

**Authors:** Ryota Gomi, Yasufumi Matsumura, Michio Tanaka, Masaru Ihara, Yoshinori Sugie, Tomonari Matsuda, Masaki Yamamoto

## Abstract

**Objectives:** Carbapenemase-producing Enterobacterales (CPE) pose serious threats to public health. Compared with clinical CPE, the genetic characteristics of environmental CPE are not well understood. This study aimed to characterize the genetic determinants of carbapenem resistance in CPE isolated from environmental waters in Japan.

**Methods:** Eighty-five water samples were collected from rivers and a lake in Japan. CPE were identified using selective media, and genome sequencing was performed for the obtained isolates (n = 21).

**Results:** Various rare/novel carbapenemases were identified: GES-5 in *Raoultella planticola* (n = 1), FRI-8 and FRI-11 in *Enterobacter* spp. (n = 8), IMI-22 and IMI-23 in *Serratia ureilytica* (n = 3), and SFC-1, SFC-2 and SFH-1 in *Serratia fonticola* (n = 9). Genomes of 11 isolates could be closed, allowing the elucidation of the genetic contexts of the carbapenemase genes. The *bla*_GES-5_ gene was located within a class 1 integron, In2071 (cassette array, *bla*_GES-5_-*aacA3-aadA16*), on a 33 kb IncP6 plasmid. The *bla*_FRI-8_ genes were carried on IncFII(Yp) plasmids ranging in size from 191 kb to 244 kb, and the *bla*_FRI-11_ genes were carried on 70 kb and 74 kb IncFII(pECLA)/IncR plasmids. The *bla*_IMI-22_ and *bla*_IMI-23_ genes were colocated on a 107 kb plasmid. The *bla*_SFC_ and *bla*_SFH-1_ genes were found on putative genomic islands inserted at tRNA-Phe genes in chromosomes.

**Conclusions:** This study revealed the presence of rare/novel carbapenemases among CPE in aquatic environments, suggesting that the environment may act as a potential reservoir of these minor carbapenemases.

## INTRODUCTION

The worldwide spread of carbapenemase-producing Enterobacterales (CPE) is a major threat to public health. CPE can be resistant to carbapenems and many other classes of commonly used drugs, which significantly limits treatment options.^1, 2^ The most clinically relevant carbapenemases detected in CPE are the ‘big five’ carbapenemases (Ambler class A KPC enzymes, class B enzymes of IMP, VIM, and NDM, and class D OXA-48-like enzymes), while Enterobacterales producing other ‘minor’ carbapenemases have also been sporadically reported. ^3^

Historically, antibiotic resistance has been considered to be a clinical problem, but a growing number of studies have suggested that the environment represents an important reservoir for antibiotic resistant bacteria and antibiotic resistance genes.^4–6^ Previous studies have reported the occurrence of CPE in environmental waters, with the aforementioned ‘big five’ carbapenemases being the predominant types.^7, 8^ Minor carbapenemases, such as GES and IMI, were also detected among environmental CPE,^9–11^ raising the possibility that environmental waters may act as reservoirs for these minor carbapenemases.

Previous studies have shown that Japanese clinical CPE usually carry *bla*_IMP_ and sporadically carry other carbapenemase genes such as *bla*_KPC_, *bla*_NDM_, and *bla*_OXA-48_.^12, 13^ However, the genetic characteristics of CPE in the natural environment, in particular in surface waters, are not well understood in Japan.^14^ Here, we isolated CPE from river and lake water in Japan and characterized the obtained CPE isolates in terms of resistance determinants, genetic contexts of the carbapenemase genes, and phylogeny.

## MATERIALS AND METHODS

### Sample collection and bacterial isolation

In total, 72 water samples were collected from 12 sites in Lake Biwa and 13 water samples were collected from 8 sites in its surrounding rivers between February and November 2020 (**Figure 1**). All samples were collected in sterile 1-L sampling bottles. Samples (100 mL to 600 mL) were processed using the membrane filter method with chromID CARBA agar (bioMérieux, Marcy-l’Étoile, France). After 20 h of incubation at 37 °C, for each sample, up to three colonies showing *Escherichia coli* profiles (pink to burgundy) or KESC (*Klebsiella, Enterobacter, Serratia, Citrobacter*) profiles (bluish green to bluish grey) with different colony morphologies were isolated. The oxidase test was performed using a cytochrome oxidase test strip (Nissui, Tokyo, Japan), and isolates with negative test results were stored at −85°C in 35% glycerol. Colonies suspected to be non-Enterobacterales species were not saved because some non-Enterobacterales species prevalent in surface waters are intrinsically resistant to carbapenems.^15, 16^

**Figure 1.**
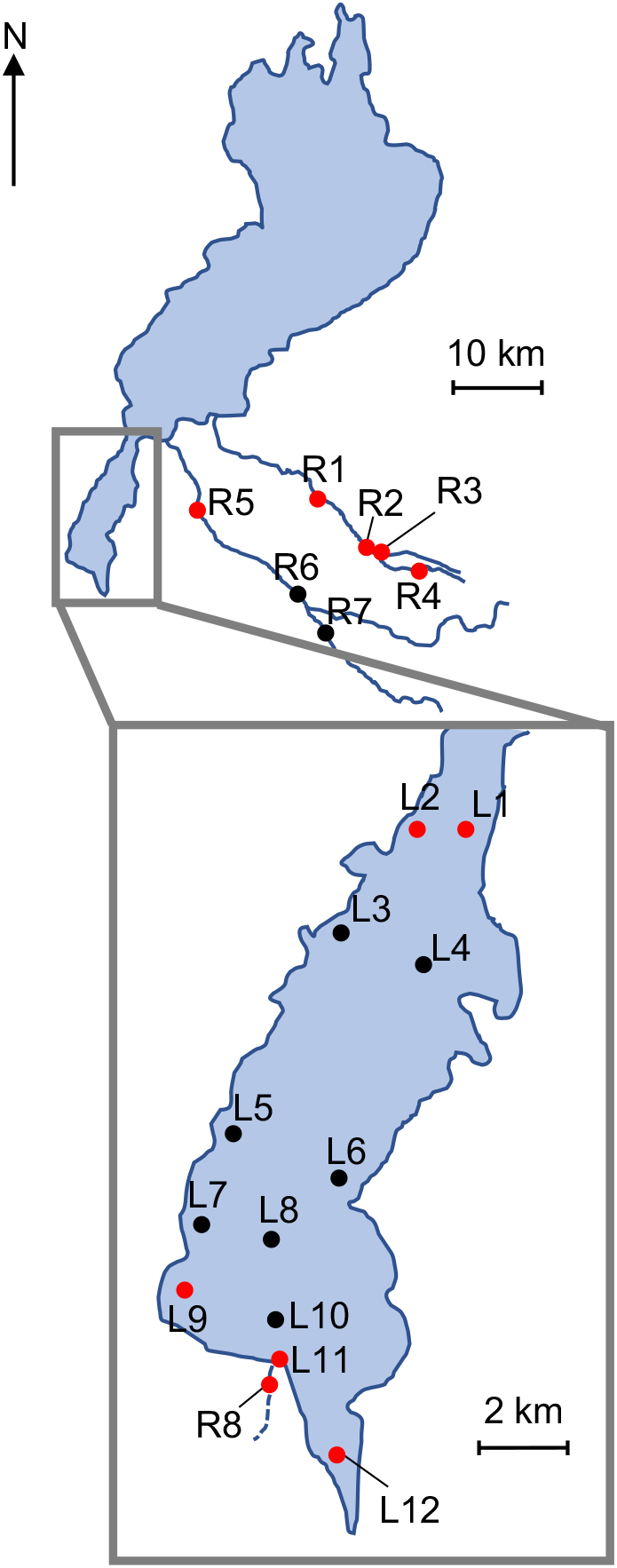
Map of sampling sites. Sampling sites in Lake Biwa are indicated with L1 to L12, and sampling sites in rivers are indicated with R1 to R8. R8 is in a small river and located near L11. Red filled circles indicate sampling points where CPE isolates were detected.

### Antibiotic susceptibility testing

Antibiotic susceptibility was assessed by microdilution using a Dry Plate Eiken (Eiken, Tokyo, Japan) using 29 antibiotics as described in the Supplementary Materials and methods. The results were interpreted using CLSI criteria.^17^ The Carba NP test was performed to test the carbapenemase activity.^18^ *Klebsiella pneumoniae* ATCC BAA-1705 was used for the positive control, and *K. pneumoniae* ATCC BAA-1706 was used for the negative control.

### Genome sequencing and assembly

Genome sequencing and assembly were performed as described in the Supplementary Materials and methods.

### Genomic analysis

Species identification was performed based on the genome sequence data, i.e., by comparing the obtained genome sequences to the genome sequences of the type strains. The average nucleotide identity (ANI) analysis was performed using OrthoANIu,^19^ and digital DNA-DNA hybridization (dDDH) analysis was performed using the GGDC website (formula 2).^20^ A ≥96% ANI value or ≥ a 70% DDH value was used for defining the species.^20, 21^ STs were assigned to *Enterobacter* isolates using the MLST scheme established previously.^22^ Whole-genome SNP-based trees were built using kSNP3.^23^

Plasmid replicons, relaxase types, mate-pair formation types, and transferability of plasmids were analysed using PlasmidFinder 2.1 and MOB-typer.^24, 25^ Annotation was performed using ISfinder,^26^ the RAST server,^27^ DFAST,^28^ ResFinder 4.1,^29^ Abricate (v1.0.1, https://github.com/tseemann/abricate), INTEGRALL,^30^ and the BLAST algorithm (https://blast.ncbi.nlm.nih.gov/Blast.cgi). Genomic islands (GIs) were detected using IslandViewer 4.^31^ New allele numbers for carbapenemase genes were assigned by NCBI through submission of sequence data according to instructions on the following website: https://www.ncbi.nlm.nih.gov/pathogens/submit-beta-lactamase/.

### Cloning experiments

Novel carbapenemase genes (*bla*_FRI-11_ in JBIWA008, *bla*_SFC-2_ in JSRIV001, *bla*_IMI-22_ and *bla*_IMI-23_ in JBIWA004) were amplified using the primers listed in **Table S1**, cloned into the pCR2.1-TOPO vector using the TOPO TA cloning kit (Thermo Fisher Scientific, Waltham, MA, USA), and transformed into *E. coli* DH5α (TaKaRa Bio, Inc., Shiga, Japan). Antibiotic susceptibilities of the recipient and transformants were assessed by microdilution as described above.

### Accession numbers

The complete genomes and sequence read files were deposited in GenBank and the NCBI SRA under BioProject PRJNA725976. The BioSample accession number for each isolate is also available in **Table 1**.

**Table 1.**
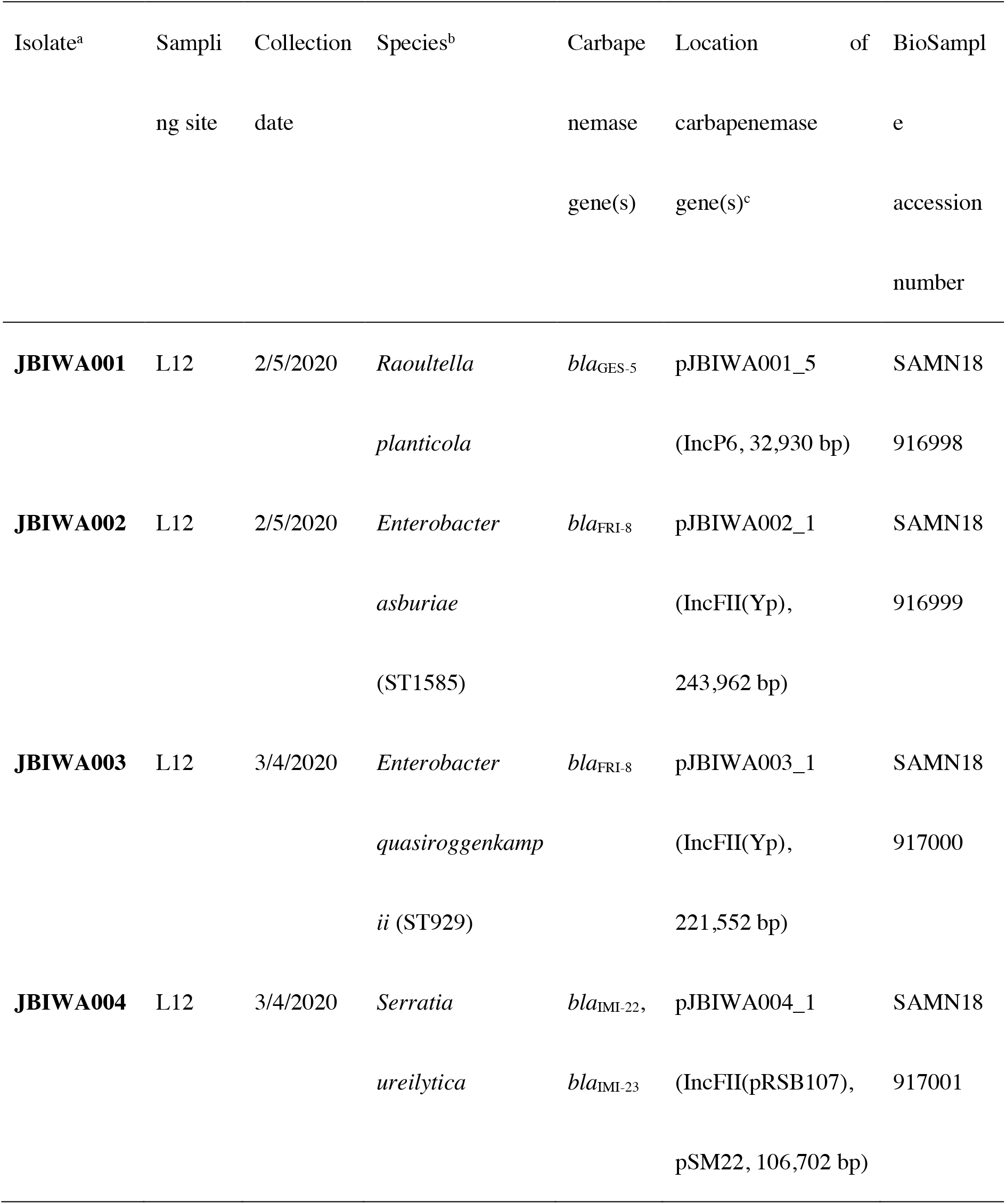
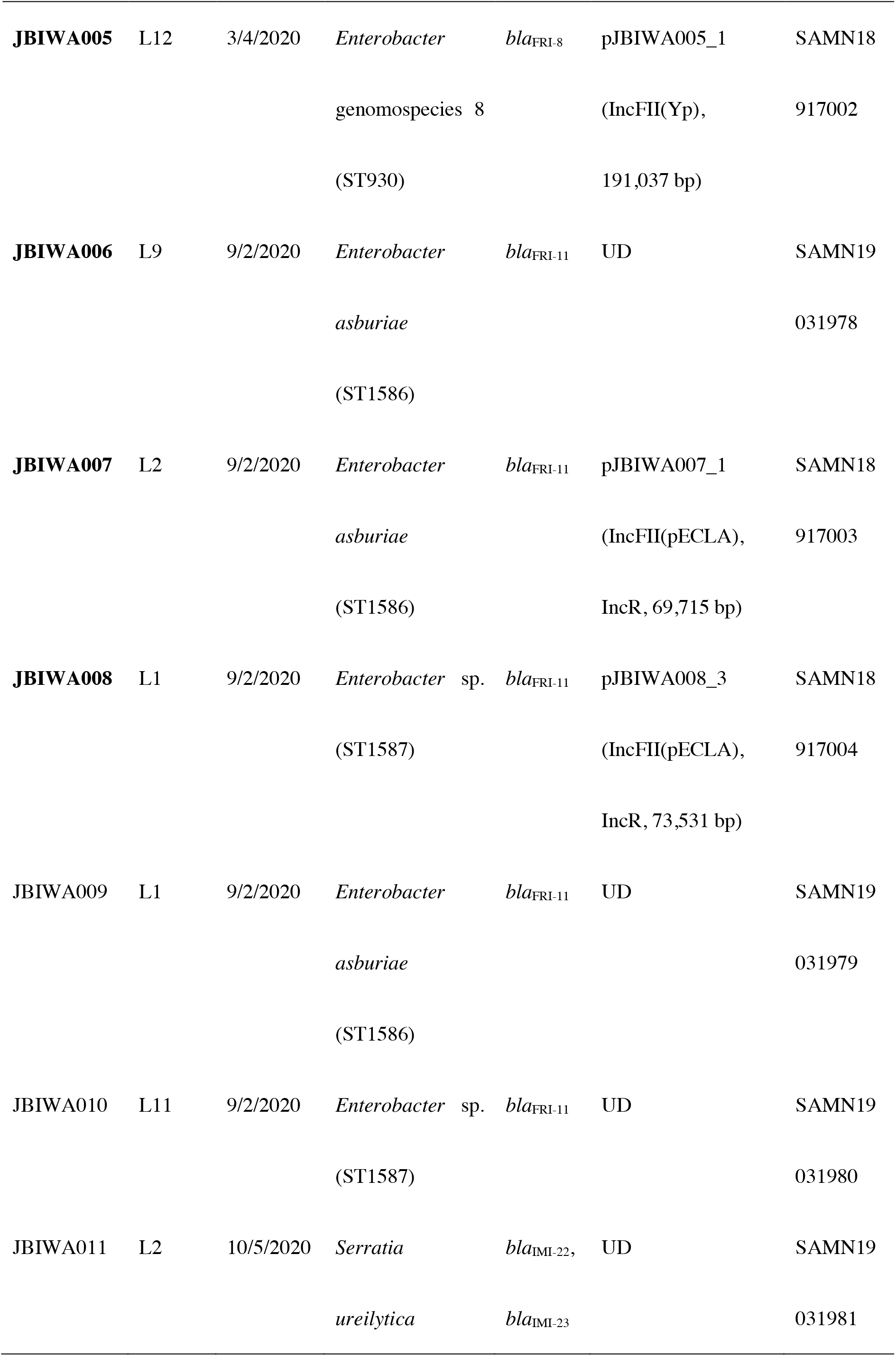
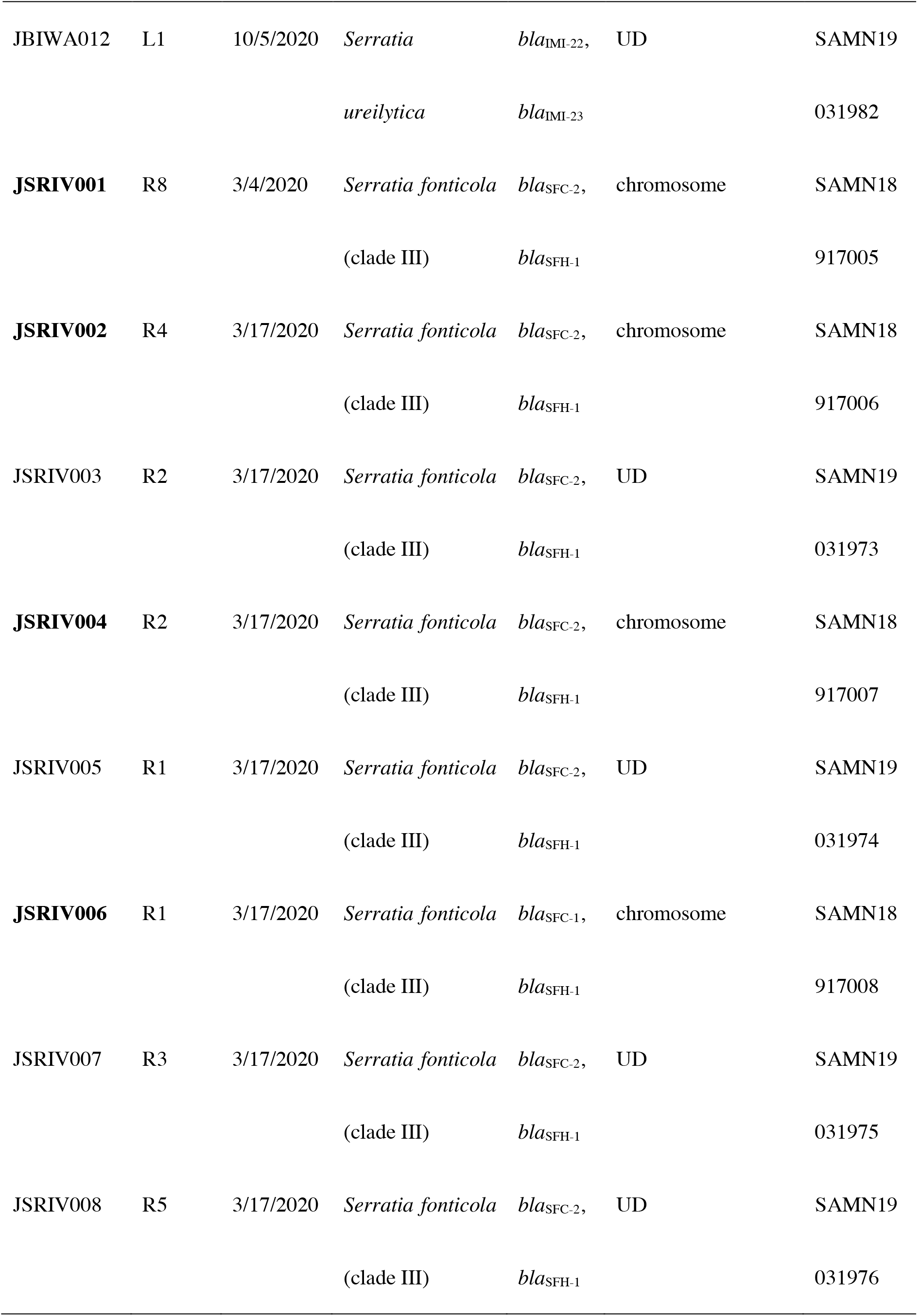
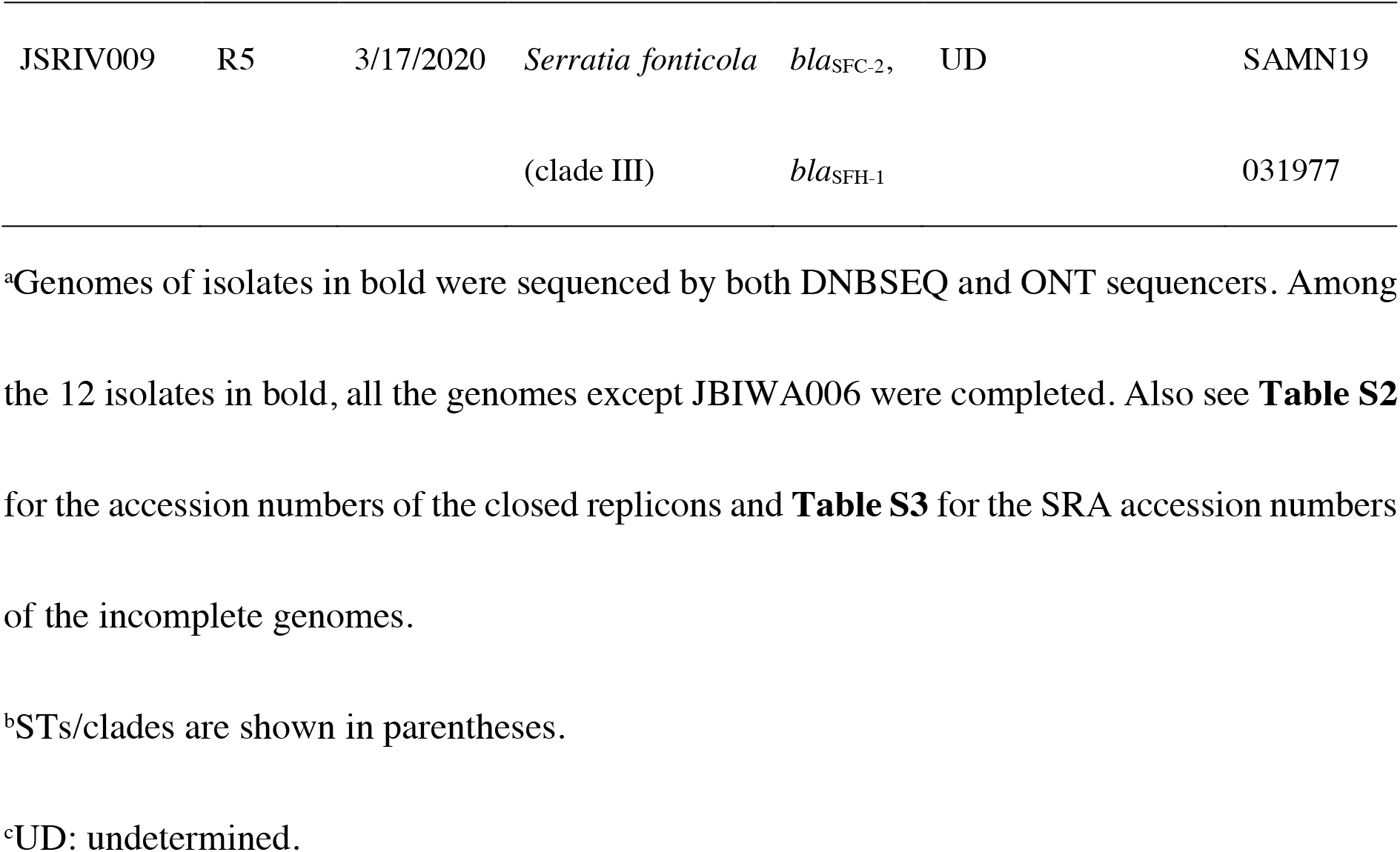
Characteristics of 21 CPE isolates recovered from surface waters in the present study.

## RESULTS AND DISCUSSION

### Detection of CPE in water samples

Twelve isolates and nine isolates were obtained from lake water samples and river water samples, respectively (**Table 1**). Lake isolates were identified as *Enterobacter* spp. (n = 8), *Serratia ureilytica* (n = 3), and *Raoultella planticola* (n = 1), and all nine river isolates were identified as *Serratia fonticola* clade III (discussed in detail below) by ANI and dDDH analyses (**Table 1**).

All 21 isolates were subjected to DNBSEQ short-read sequencing, and 12 isolates were also subjected to Oxford Nanopore Technologies (ONT) long-read sequencing. Among the 12 isolates with both short-read and long-read data, all genomes except for the genome of JBIWA006 were completed by either Unicycler^32^ or Flye.^33^ The completed genomes contained one to 12 plasmids (see **Table S2** for details). The draft genome assemblies of the remaining isolates had 28 to 217 contigs and 189,689 bp to 2,472,998 bp N50 values after discarding contigs shorter than 100 bp (see **Table S3** for details).

All the carbapenemase genes detected in this study were found to encode minor carbapenemases, namely, *bla*_GES-5_, *bla*_FRI_, *bla*_IMI_, *bla*_SFC_, which all encode Ambler class A and functional group 2f enzymes, and *bla*_SFH-1_, which encodes an Ambler subclass B2 and functional group 3b enzyme (GES: Guiana Extended-Spectrum β-lactamase, FRI: French imipenemase, IMI: Imipenem-hydrolysing β-lactamase, SFC: *Serratia fonticola* carbapenemase, SFH: *Serratia fonticola* carbapenem hydrolase).^34, 35^ Briefly, *bla*_GES-5_ was detected in a *R.planticola* isolate, *bla*_FRI-8_ and *bla*_FRI-11_ were detected in *Enterobacter* isolates, *bla*_IMI-22_ and *bla*_IMI-23_ were detected in *S. ureilytica* isolates, and *bla*_SFC-1_, *bla*_SFC-2_ and *bla*_SFH-1_ were detected in *S. fonticola* clade III isolates (**Table 1**). *bla*_FRI-11_, *bla*_IMI-22_, bla_IMI-23_, and *bla*_SFC-2_ are the genes encoding novel carbapenemase variants, and their protein accession numbers are as follows: FRI-11 (WP_223174987.1), IMI-22 (WP_223174988.1), IMI-23 (WP_223525081.1), and SFC-2 (WP_223174986.1).

The carbapenemases and bacterial species detected in this study are different from those usually observed in clinical settings. One possible reason for this is that these CPE isolates are better adapted to the environment than those usually observed in clinical settings. There are three municipal wastewater treatment plants (WWTPs) near the study area, and the effluents (although biologically treated) from those WWTPs could be the potential sources of CPE detected in the present study. Moreover, insufficiently treated wastewater could be released from those WWTPs on rainy days, which could also have contributed to the dissemination of CPE. The actual human health risks associated with the presence of these CPE isolates in surface waters remain unclear. However, there seem to be some overlaps between the CPE detected in this study and CPE isolated in clinical settings in terms of species and carbapenemase types. For example, FRI-producing *Enterobacter* spp. have been previously reported in patients at hospitals.^36, 37^ On the other hand, *bla*_SFC_ and *bla*_SFH_, which were prevalent among the river *Serratia* isolates, have been described only once in environmental *S. fonticola* and not in clinical settings.^3, 38–40^ Further studies are needed to elucidate the health risks posed by these CPE in surface waters.

### Susceptibility testing and cloning experiments

Phenotypic resistance was determined by microdilution. All the isolates were nonsusceptible to doripenem and meropenem, and all but GES-5-producing *R. planticola* (JBIWA001) were nonsusceptible to imipenem (**Table S4**). The Carba NP test confirmed carbapenemase activity in all isolates carrying *bla*_SFC_ and *bla*_SFH-1_, whereas isolates carrying *bla*_GES-5_, *bla*_FRI-8_, and *bla*_IMI_ were negative by the test. Results of the Carba NP test for the *bla*_FRI-11_-carrying isolates were inconsistent (two were positive, one was negative, and two were indeterminate). Previous studies reported that Enterobacterales harbouring carbapenemases such as *bla*_GES-5_ and *bla*_IMI_ can be falsely negative in the Carba NP test.^41–43^ Novel carbapenemase genes were cloned into the pCR2.1-TOPO vector and transformed into *E. coli* DH5α. An increase in MICs of carbapenems was detected in *E. coli* DH5α carrying the recombinant plasmids pCR2.1-TOPO-*bla*_SFC-2_ and pCR2.1-TOPO-*bla*_IMI-23_ (**Table S5**). However, no detectable increase in MICs of carbapenems was observed for *E. coli* DH5α carrying the recombinant plasmid pCR2.1-TOPO-*bla*_FRI-11_ or pCR2.1-TOPO-*bla*_IMI-22_ under the conditions in the present study, which indicates that FRI-11 and IMI-22 are enzymes with low carbapenemase activity.

### Genomic analysis of a *Raoultella planticola* isolate

River isolate JBIWA001 showed 99.15% ANI and 93.90% dDDH values with the type strain of *R. planticola*. The assembly resulted in a chromosome (5,626,881 bp) and 12 plasmids ranging from 1,443 bp to 136,413 bp (see **Table S2** for details). The *bla*_GES-5_ gene was on a 32,930 bp plasmid, pJBIWA001_5, with an IncP6 replicon and a MOBP-type relaxase. *bla*_GES-5_ was found to be part of a novel class 1 integron, In2071. In2071 contains a *bla*_GES-5_-*aacA3-aadA16* cassette array, followed by the 3’-conserved segment (3’-CS) (*qacEΔ1, sul1*, orf5, orf6), IS*1326, tniBΔ2, tniA*, and IRt (inverted repeat at *tni* end) (see Supplementary Results and **Figure S1** for comparative analysis of pJBIWA001_5 and a closely related plasmid).

### Genomic analysis of *Enterobacter* isolates

The *Enterobacter* genus comprises 22 species and 22 additional genomospecies.^44^ The lake *Enterobacter* isolates in the present study were identified as *E. asburiae* (n = 4), *E. quasiroggenkampii* (n = 1), and *Enterobacter* genomospecies 8 (n = 1) based on the ANI and dDDH analyses (also see **Figure S2** for the *Enterobacter* phylogenetic tree). The remaining two isolates, JBIWA008 and JBIWA010, showed < 96% ANI values and < 70% dDDH values against all type strains and reference strains (closest to *E. asburiae*, 95.2% ANI and 63.0% dDDH). It should be noted that although we adopted a 96% ANI cutoff value, which is the cutoff value used in a recent study defining *Enterobacter* species,^45^ a 95% ANI cutoff value is also used for defining the species and 95-96% ANI values can be an inconclusive zone.^45, 46^ However, dDDH values were lower than the 70% cutoff and thus these two isolates were considered to belong to a novel genomospecies. MLST assigned two novel STs (ST1585 and ST1586) to *E. asburiae* isolates, ST929 to an *E. quasiroggenkampii* isolate, ST930 to an *Enterobacter* genomospecies 8 isolate, and a novel ST (ST1587) to two isolates belonging to the novel genomospecies. Interestingly, *Enterobacter* ST929 and ST930 were first reported in our previous study analysing wastewater CPE isolates in the same region.^47^ As discussed above, the potential sources of these *Enterobacter* isolates could be the effluents from WWTPs.

We could determine the complete sequences of three *bla*_FRI-8_-carrying plasmids, namely pJBIWA002_1 (243,962 bp) from *E. asburiae* isolate JBIWA002, pJBIWA003_1 (221,552 bp) from *E. quasiroggenkampii* isolate JBIWA003, and pJBIWA005_1 (191,037 bp) from *Enterobacter* genomospecies 8 isolate JBIWA005. A BLASTN search against the GenBank database using *bla*_FRI-8_ as a query sequence identified one *bla*_FRI-8_-carrying plasmid, *Enterobacter* sp. 18A13 plasmid pECC18A13-1 (150,509 bp), in which *bla*_FRI-8_ was first detected.^37^ The BLASTN search also identified the *bla*_FLC-1_-carrying plasmid p3442-FLC-1 (93,658 bp) from the *Enterobacter cloacae* complex strain E3442.^48^ FLC-1 (FRI-like carbapenemase-1) shares 99.66% amino acid identity with FRI-8 and was suggested that it be reclassified as an FRI variant.^3^ All five plasmids (pJBIWA002_1, pJBIWA003_1, pJBIWA005_1, pECC18A13-1, p3442-FLC-1) carried an IncFII(Yp)-like replicon and were typed as Y10:A-:B-by plasmid MLST. All five plasmids also carried a MOBP-type relaxase, though pECC18A13-1 also carried a MOBF-type relaxase. Although the five plasmids carried some sequences in common, insertions/deletions contributed to variable region diversification (**Figure 2A**). Analysis of genetic contexts of *bla*_FRI-8_ and *bla*_FLC-1_ revealed the presence of an IS*Peat2*-like IS downstream of *bla*_FRI-8_/*bla*_FLC-1_ and a LysR-type transcriptional regulator upstream of *bla*_FRI-8_/*bla*_FLC-1_ (**Figure 2B**). A truncated IS*Eae1*-like element disrupted the IS*Peat2*-like IS in pJBIWA003_1, changing the downstream structure in this plasmid. RAST identified a coding sequence encoding a hypothetical protein between *bla*_FRI-8_/*bla*_FLC-1_ and a LysR-type transcriptional regulator. BLASTN analysis showed this coding sequence to be absent near the other *bla*_FRI_ genes.

**Figure 2.**
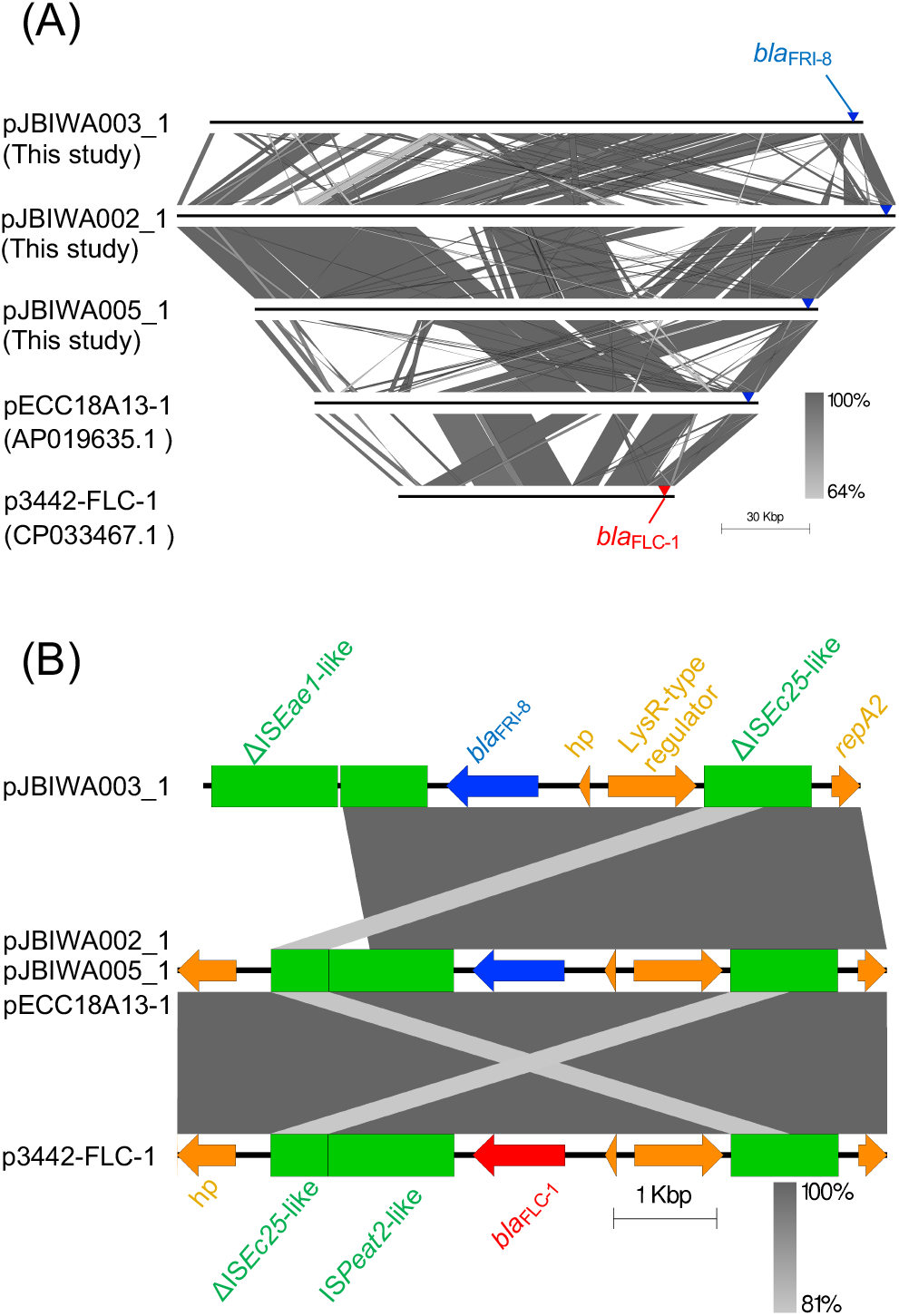
(A) Alignment of plasmids pJBIWA003_1, pJBIWA002_1, pJBIWA005_1, pECC18A13-1, and p3442-FLC-1. Positions of *bla*_FRI-8_ and *bla*_FLC-1_ are indicated by blue and red triangles, respectively. (B) Genetic contexts of *bla*_FRI-8_ and *bla*_FLC-1_. Blue arrows indicate *bla*_FRI-8_, a red arrow indicates *bla*_FLC-1_, green boxes indicate insertion sequences, and orange arrows indicate other coding sequences. hp: hypothetical protein. The alignments were generated with Easyfig.^54^

We were able to determine the complete sequences of two *bla*_FRI-11_-carrying plasmids, namely pJBIWA007_1 (69,715 bp) from *E. asburiae* isolate JBIWA007 and pJBIWA008_3 (73,531 bp) from *Enterobacter* sp. isolate JBIWA008. Online BLASTN analysis identified a 108,672 bp plasmid, pJF-587, from *E. asburiae*,^49^ which showed >90% coverage to pJBIWA007_1 and pJBIWA008_3. pJF-587 carried *bla*_FRI-2_, which shares 98.5% nucleotide identity with *bla*_FRI-11_. All three plasmids carried IncFII(pECLA) and IncR replicons. Although pJF-587 contained a MOBF-type relaxase and was predicted to be conjugative by MOB-typer, pJBIWA007_1 and pJBIWA008_3 did not contain a relaxase and were predicted to be non-mobilizable. Alignment of the three plasmids revealed conjugation genes present on pJF-587 were deleted in pJBIWA007_1 and pJBIWA008_3 (**Figure 3A**). The genetic contexts of *bla*_FRI-11_/*bla*_FRI-2_ were slightly different among the three plasmids. A DNA segment containing *ltrA* was present in pJBIWA008_3, while this region was absent in pBIWA007_1 (**Figure 3B**). In pJF-587, this segment was inverted compared to pJBIWA008_3, and an IS*Sm3*-like element was absent. Although complete plasmid sequences were not available for JBIWA006, JBIWA009, and JBIWA010, we could determine the genetic contexts of *bla*_FRI-11_ in these isolates using short-read assemblies: a pBIWA007_1-type genetic context was detected in JBIWA006 and JBIWA009, while a pJBIWA008_3-type genetic context was detected in JBIWA010.

**Figure 3.**
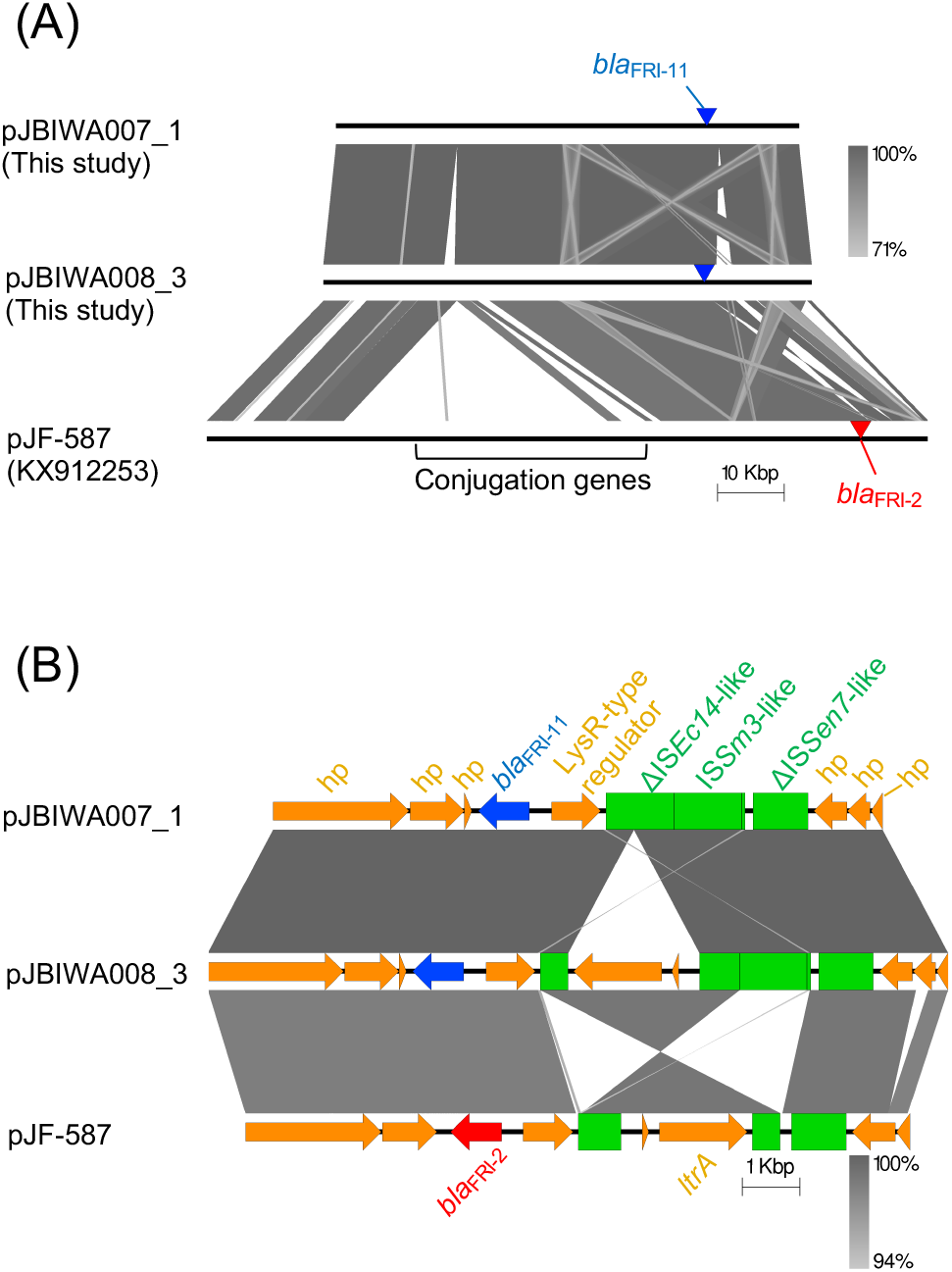
(A) Alignment of pJBIWA007_1, pJBIWA008_3, and pJF-587. Positions of *bla*_FRI-11_ and *bla*_FRI-2_ are indicated by blue and red triangles, respectively. (B) Genetic contexts of *bla*_FRI-11_ and *bla*_FRI-2_. Blue arrows indicate *bla*_FRI-11_, a red arrow indicates *bla*_FRI-2_, green boxes indicate insertion sequences, and orange arrows indicate other coding sequences.

### Genomic analysis of *Serratia* isolates

Although three *Serratia* isolates recovered from the lake could be classified as *S. ureilytica*, nine *Serratia* isolates from rivers could not be classified based on the ANI and dDDH analyses. The pairwise ANI values and dDDH values among the nine river isolates were all above 96% and 70%, respectively, indicating the river isolates belong to a single species. The river isolates seem to be related to *S. fonticola* (**Figure S3**), but the ANI and dDDH values between the river isolates and the type strain of *S. fonticola* ranged from 95.5% to 95.7% and 65.1% to 65.4%, respectively, which are lower than the 96% ANI and 70% dDDH cutoff values. These results suggest the 9 river isolates could belong to a novel species whose closest relative is *S. fonticola*. It should be noted that genome sequences for type strains were not available for some *Serratia* species (*Serratia aquatilis*, *Serratia bozhouensis*, *Serratia entomophila*, *Serratia myotis*, and *Serratia profundus*), and ANI and dDDH analyses could not be performed for these species. However, 16S rRNA gene sequences were available for them and all showed < 98.7% identity with the 16S rRNA gene sequences of the river isolates, which implies they belong to different species.^46^

In order to gain further insight into the phylogenetic characteristics of river *Serratia* isolates, we downloaded RefSeq assemblies labelled *S. fonticola* and built a phylogenetic tree (**Figure 4**, also see **Figure S4** for phylogenetic characteristics of these strains in relation to other *Serratia* species). Interestingly, three clades were identified in the phylogenetic tree, namely clade I (containing the genomes of type strains of *S. fonticola*), clade II, and clade III (containing the genomes of river isolates). Mean within-clade ANI values were > 98% for all clades, and within-clade pairwise ANI values always exceeded 96%. On the other hand, mean between-clade ANI values were < 96%, and between-clade pairwise ANI values were always < 96% (**Table S6**). As there are limitations in the use of the website-based GGDC dDDH calculation tool to compare large numbers of genomes, we performed dDDH analysis only for comparisons including the type strain from clade I or the reference strains from clade II and clade III (**Table S7**). The results show that within-clade pairwise dDDH values were always above 70% while between-clade pairwise dDDH values were always below 70%. These results suggest clades I-III could represent different species, although further studies are needed to confirm this.

**Figure 4.**
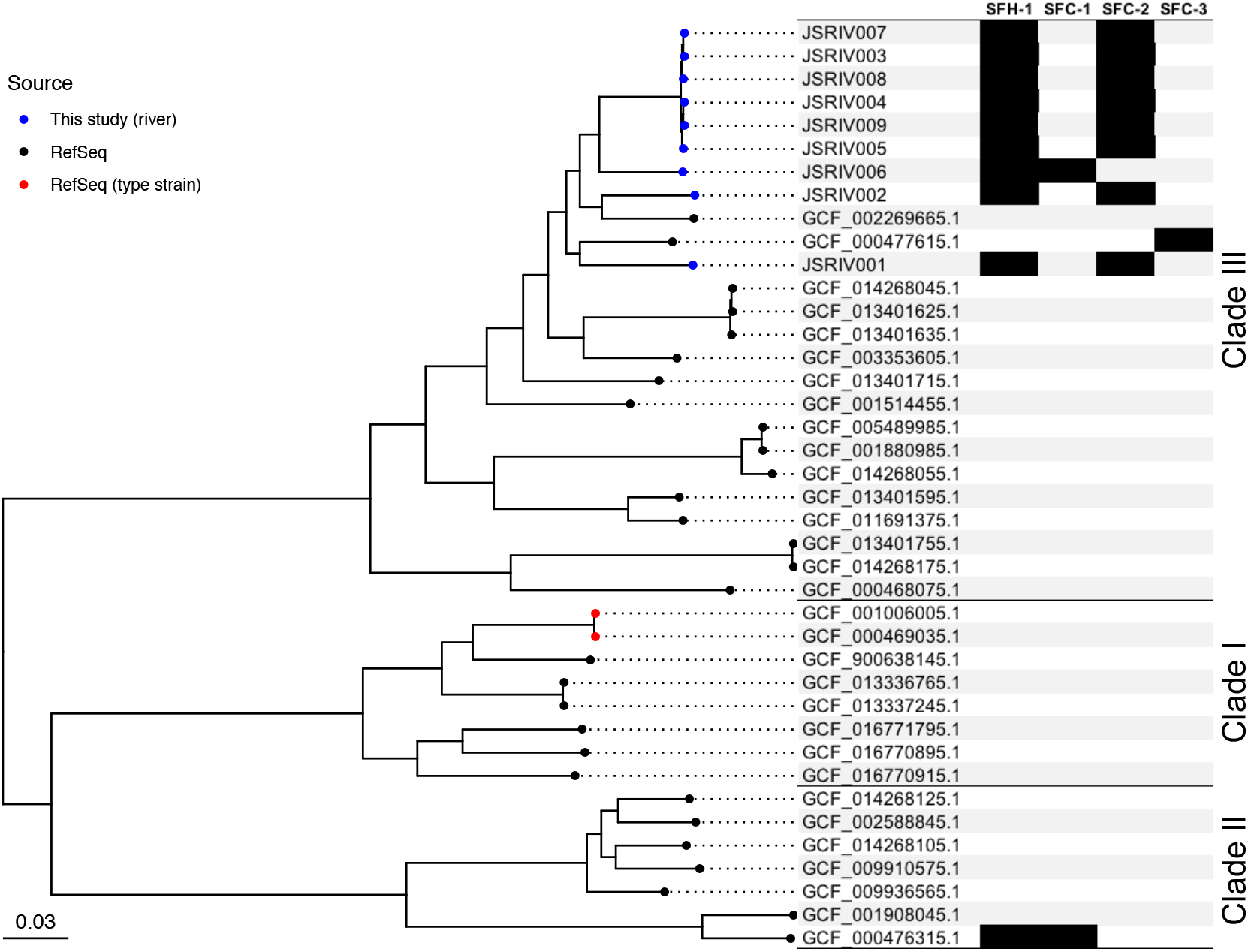
Phylogenetic tree constructed using RefSeq assemblies labelled *S. fonticola* and assemblies of nine river *Serratia* isolates. The RefSeq assemblies were downloaded in March 2021. The tree was constructed based on 177,349 SNPs occurring in at least 80% of the genomes. Four RefSeq assemblies (GCF_006714955.1, GCF_006715025.1, GCF_007827525.1, GCF_014205655.1) were removed because these assemblies showed < 85% ANI values with the type strain of *S. fonticola*. The mid-point rooted tree was visualized with ggtree.^55^ The black blocks represent the presence of a carbapenemase gene.

River *Serratia* isolates carried two types of carbapenemase genes, *bla*_SFH_ and *bla*_SFC_. *bla*_SFH-1_ and *bla*_SFC-1_ have been described only once in *S. fonticola* clade II strain UTAD54 (GCF_000476315.1) from Portugal.^3, 38–40^ Although no variants were reported previously for either of these two carbapenemases, online BLASTN analysis revealed that *S. fonticola* clade III strain AU-AP2C (GCF_000477615.1)^50^ carried a variant of *bla*_SFC_ (the product shows 99.4% amino acid identity to SFC-1). All river isolates except JSRIV006 carried *bla*_SFH-1_ and a variant of *bla*_SFC_ that is different from *bla*_SFC-1_ and *bla*_SFC_ of AU-AP2C (the product shows 99.4% amino acid identities to SFC-1 and SFC of AU-AP2C). Novel alleles SFC-2 and SFC-3 were assigned to SFC enzymes in river isolates and AU-AP2C, respectively. JSRIV006, in contrast, carried *bla*_SFH-1_ and *bla*_SFC-1_, though a synonymous mutation was detected in *bla*_SFH-1_ when compared with *bla*_SFH-1_ in UTAD54. Although detailed genetic contexts of the *bla*_SFH_ and *bla*_SFC_ genes are not known, these genes were previously shown to be located on the chromosome of UTAD54 by probe hybridization.^38, 39^ The *bla*_SFH-1_ and *bla*_SFC_ genes in the river isolates sequenced to closure in this study (JSRIV001, JSRIV002, JSRIV004, JSRIV006) were also located on chromosomes and situated close to each other (separated by 76 kbp to 131 kbp). Given the sporadic phylogenetic distribution (**Figure 4**), the two genes were suspected to be located on GIs.^51^ Comparative analysis of the chromosomes of four river isolates and the chromosome of *S. fonticola* clade III strain GS2 (GCF_001514455.1) revealed that the two genes were located on putative GIs inserted at tRNA-Phe genes. (**Figure 5A**). Importantly, IslandViewer 4 predicted almost the same regions as potential GIs (**Figure S5**). Perfect (5’-TCGATTCCGAGTCCGGGCACCA-3’, detected in JSRIV004) or imperfect (5’-TCGATTCCGAGTCC-GGCACCA-3’, detected in JSRIV001, JSRIV002, and JSRIV006) repeat sequences corresponding to the 3’-end of the tRNA-Phe gene were detected at the right ends of the GIs. Some previous studies also reported imperfect repeats at the right ends of GIs.^52, 53^ An integrase gene was detected ~200 bp downstream of the tRNA-Phe gene in each GI (**Figure 5B**), which indicates that the GIs were integrated at tRNA-Phe genes by site-specific recombination mediated by these integrases. Genetic contexts of *bla*_SFH-1_ were highly conserved and the *bla*_SFH-1_ gene was surrounded by a ΔIS*Ec18*-like element and an IS*1N*-like element in all four completed chromosomes (**Figure 5B**). Although the complete genome was not available for strain UTAD54, analysis of a contig carrying *bla*_SFH-1_ revealed that the gene was embedded within the same genetic context and also carried by a putative GI which was inserted at the tRNA-Phe gene (**Figure 5B**). In draft assemblies of JSRIV003, JSRIV005, JSRIV007, JSRIV008 and JSRIV009, contigs were broken near *bla*_SFH-1_ and genetic contexts of *bla*_SFH-1_ could not be determined. Genetic surroundings of *bla*_SFC_ were also highly conserved, though variations were observed in regions >3,000 bp away from *bla*_SFC_ (**Figure 5C**). *bla*_SFC_ was surrounded by genes such as *smvA* (encoding a methyl viologen resistance protein), *acrR* (encoding a transcriptional regulator), and genes encoding hypothetical proteins. The genetic surroundings of *bla*_SFC_ (<3,000 bp from *bla*_SFC_) were also conserved in draft assemblies of JSRIV003, JSRIV005, JSRIV007, JSRIV008 and JSRIV009. Although *bla*_SFH_ and *bla*_SFC_ seem to be mobilized together within the same GIs, strain AU-AP2C carried only *bla*_SFC-3_ and lacked *bla*_SFH_. A contig carrying *bla*_SFC_-3 in the draft genome assembly of AU-AP2C was long enough for the comparative analysis. In AU-AP2C, a putative GI was inserted at the tRNA-Phe gene, and *bla*_SFC-3_ was also carried on this putative GI (**Figure S6A**). However, the region containing the integrase gene and *bla*_SFH-1_ was replaced by another segment as shown in **Figure S6B**.

**Figure 5.**
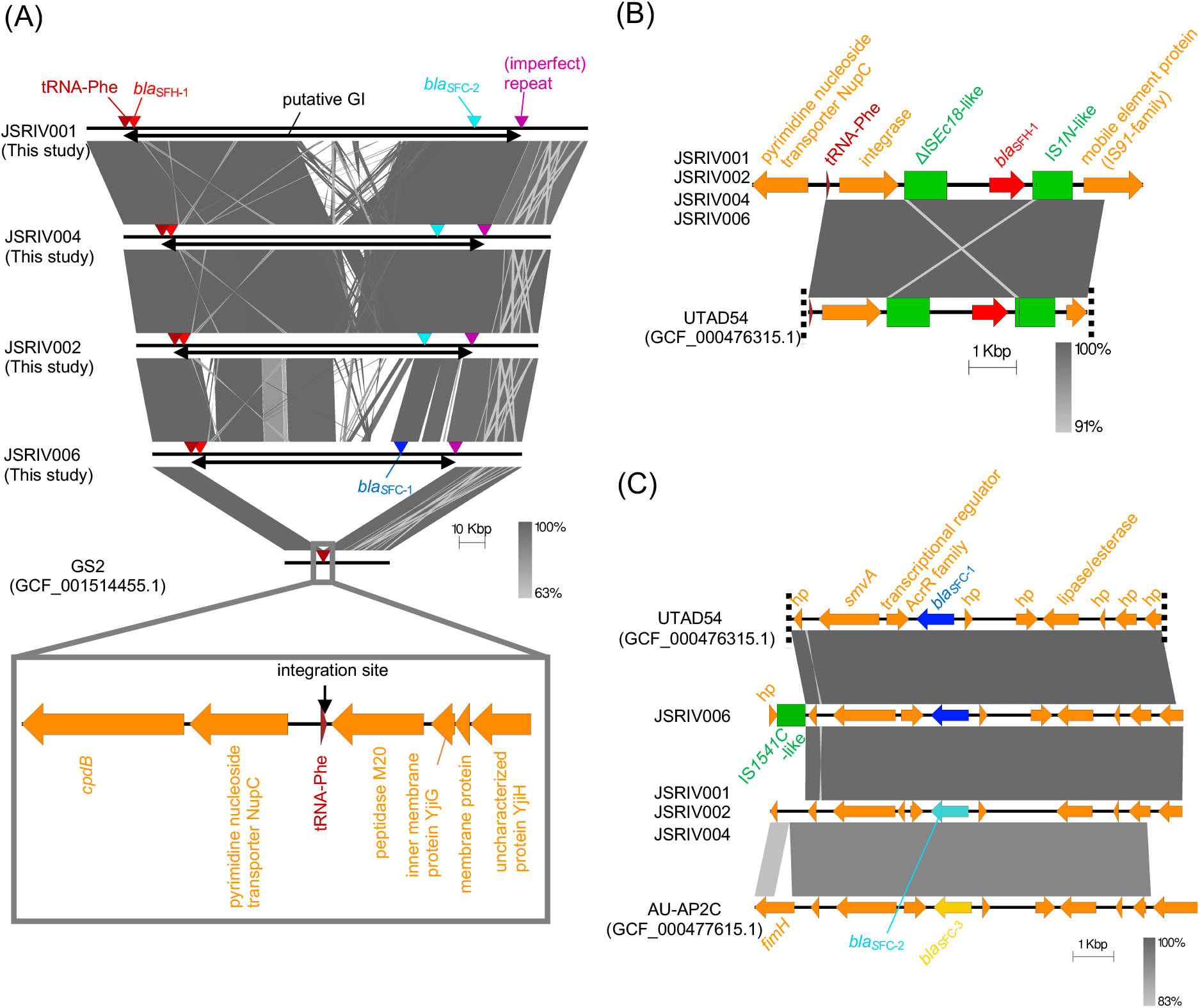
(A) Alignment of the putative GIs and the surrounding regions of river *Serratia* isolates sequenced to closure and the corresponding chromosomal region of *S. fonticola* clade III strain GS2. Positions of the *bla*_SFC-1_, *bla*_SFC-2_, *bla*_SFH-1_, and tRNA-Phe genes, and (imperfect) repeats are indicated by blue, light blue, red, brown, and purple triangles, respectively. Double-headed arrows indicate genomic regions corresponding to GIs. Genetic contexts of (B) *bla*_SFH_ and (C) *bla*_SFC_. Red arrows indicate *bla*_SFH-1_, blue arrows indicate *bla*_SFC-1_, a light blue arrow indicates *bla*_SFC-2_, a yellow arrow indicates *bla*_SFC-3_, green boxes indicate insertion sequences, and orange arrows indicate other coding sequences. Dashed lines indicate contig breaks.

Three *S. ureilytica* isolates from the lake, JBIWA004, JBIWA011 and JBIWA012, carried *bla*_IMI-22_ and *bla*_IMI-23_. Among the three isolates, the genome of JBIWA004 was sequenced to closure. The two *bla*_IMI_ genes in JBIWA004 were located on a 106,702 bp plasmid, named pJBIWA004_1, with IncFII(pRSB107) and pSM22 replicons. Online BLASTN analysis revealed the plasmid p2-125 from *Serratia liquefaciens* showed the highest coverage (55%) and also showed the high percent identity (96%) to pJBIWA004_1, while other plasmids showed <30% coverage. Alignment of pJBIWA004_1 and p2-125 revealed that the two plasmids shared conjugation regions, but the region containing the *bla*_IMI_ genes was only present in pJBIWA004_1 (**Figure 6A**). Comparison of the genetic contexts of the two *bla*_IMI_ genes and that of *bla*_IMI-1_ in *E. cloacae* complex strain N11-1168 showed that the regulator *imiR* gene and a gene encoding a hypothetical protein, which are usually present upstream and downstream of *bla*_IMI_ genes, were truncated or deleted in pJBIWA004_1 (**Figure 6B**). *bla*_IMI-22_ and *bla*_IMI-23_ were surrounded by remnants of ISs, which might have contributed to their mobilization. Although only draft genome assemblies were available for JBIWA011 and JBIWA012, the *bla*_IMI-22_ and *bla*_IMI-23_ genes were located on a single contig in each assembly. Moreover, those contigs showed >94% coverage and >99.9% identity to pJBIWA004_1, indicating that JBIWA011 and JBIWA012 also carried *bla*_IMI-22_ and *bla*_IMI-23_ on pJBIWA004_1-like plasmids.

**Figure 6.**
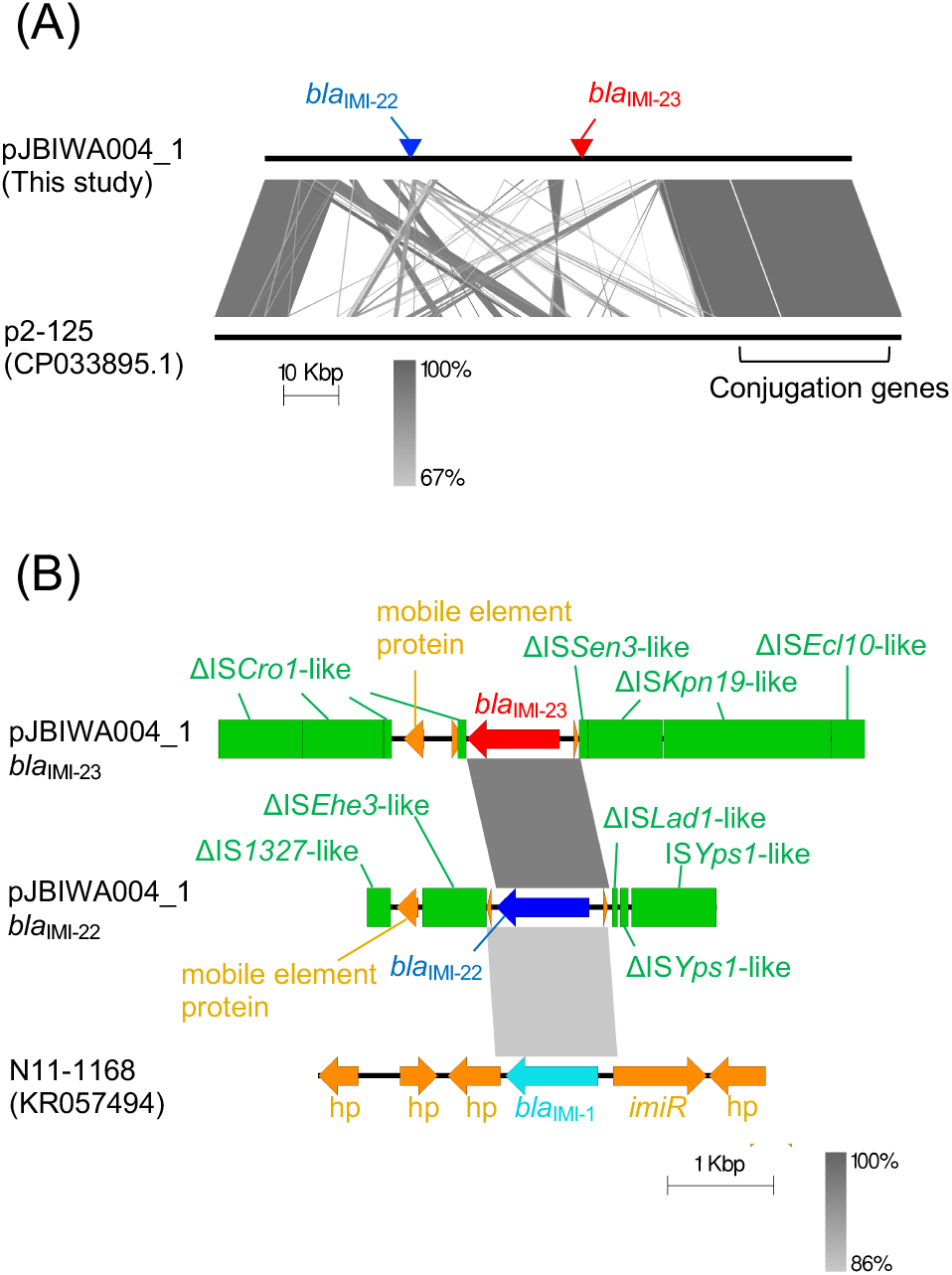
(A) Alignment of pJBIWA004_1 and p2-125. Positions of *bla*_IMI-22_ and *bla*_IMI-23_ are indicated by blue and red triangles, respectively. (B) Genetic contexts of *bla*_IMI_ genes. A blue arrow indicates *bla*_IMI-22_, a red arrow indicates *bla*_IMI-23_, a light blue arrow indicates *bla*_IMI-1_, green boxes indicate insertion sequences, and orange arrows indicate other coding sequences.

### Conclusions

Here, we identified the prevalence of minor carbapenemases among environmental CPE isolates in Japan. Of note, *bla*_SFH_ and *bla*_SFC_, which have been described only in one *S. fonticola* isolate in Portugal, were prevalent among river *S. fonticola* isolates obtained from different sampling sites. The results of this study indicate aquatic environments as potential reservoirs of minor carbapenemases and highlight the need to continuously monitor antibiotic resistance in the environment.

## Supporting information

Supplementary Materials and methods, Supplementary Results, Supplemental Tables (S1, S5-S7), and Supplementary Figures (S1-S6)

## Acknowledgements

Computations were partially performed on the NIG supercomputer at ROIS National Institute of Genetics.

## Funding

This work was supported by the Japan Society for the Promotion of Science KAKENHI (grant number JP19K20461) and the Environment Research and Technology Development Fund (JPMEERF20205006) of the Ministry of the Environment Japan.

## Transparency declarations

None to declare.

## Notes

### Competing Interest Statement

The authors have declared no competing interest.

## References

1. Tzouvelekis LS, Markogiannakis A, Piperaki E et al. Treating infections caused by carbapenemase-producing Enterobacteriaceae. Clin Microbiol Infect 2014; 20: 862–72.

2. Doi Y, Paterson DL. Carbapenemase-producing *Enterobacteriaceae*. Semin Respir Crit Care Med 2015; 36: 74–84.

3. Bonnin RA, Jousset AB, Emeraud C et al. Genetic Diversity, Biochemical Properties, and Detection Methods of Minor Carbapenemases in Enterobacterales. Front Med (Lausanne) 2020; 7: 616490.

4. Marti E, Variatza E, Balcazar JL. The role of aquatic ecosystems as reservoirs of antibiotic resistance. Trends Microbiol 2014; 22: 36–41.

5. Zainab SM, Junaid M, Xu N et al. Antibiotics and antibiotic resistant genes (ARGs) in groundwater: A global review on dissemination, sources, interactions, environmental and human health risks. Water Res 2020; 187: 116455.

6. Berglund B. Environmental dissemination of antibiotic resistance genes and correlation to anthropogenic contamination with antibiotics. Infect Ecol Epidemiol 2015; 5: 28564.

7. Bleichenbacher S, Stevens MJA, Zurfluh K et al. Environmental dissemination of carbapenemase-producing Enterobacteriaceae in rivers in Switzerland. Environ Pollut 2020; 265: 115081.

8. Tafoukt R, Touati A, Leangapichart T et al. Characterization of OXA-48-like-producing *Enterobacteriaceae* isolated from river water in Algeria. Water Res 2017; 120: 185–9.

9. Suzuki Y, Nazareno PJ, Nakano R et al. Environmental Presence and Genetic Characteristics of Carbapenemase-Producing *Enterobacteriaceae* from Hospital Sewage and River Water in the Philippines. Appl Environ Microbiol 2020; 86: e01906–19.

10. Teixeira P, Tacao M, Pureza L et al. Occurrence of carbapenemase-producing *Enterobacteriaceae* in a Portuguese river: *bla*_NDM_, *bla*_KPC_ and *bla*_GES_ among the detected genes. Environ Pollut 2020; 260: 113913.

11. Mathys DA, Mollenkopf DF, Feicht SM et al. Carbapenemase-producing *Enterobacteriaceae* and *Aeromonas* spp. present in wastewater treatment plant effluent and nearby surface waters in the US. PLoS One 2019; 14: e0218650.

12. Ohno Y, Nakamura A, Hashimoto E et al. Molecular epidemiology of carbapenemase-producing Enterobacteriaceae in a primary care hospital in Japan, 2010-2013. J Infect Chemother 2017; 23: 224–9.

13. Eda R, Nakamura M, Takayama Y et al. Trends and molecular characteristics of carbapenemase-producing Enterobacteriaceae in Japanese hospital from 2006 to 2015. J Infect Chemother 2020; 26: 667–71.

14. Cherak Z, Loucif L, Moussi A et al. Carbapenemase-producing Gram-negative bacteria in aquatic environments: a review. J Glob Antimicrob Resist 2021; 25: 287–309.

15. Tacao M, Correia A, Henriques IS. Low Prevalence of Carbapenem-Resistant Bacteria in River Water: Resistance Is Mostly Related to Intrinsic Mechanisms. Microb Drug Resist 2015; 21: 497–506.

16. Harmon DE, Miranda OA, McCarley A et al. Prevalence and characterization of carbapenem-resistant bacteria in water bodies in the Los Angeles-Southern California area. Microbiologyopen 2019; 8: e00692.

17. CLSI. Performance Standards for Antimicrobial Susceptibility Testing—Thirtieth Edition: M100. 2020.

18. Nordmann P, Poirel L, Dortet L. Rapid detection of carbapenemase-producing *Enterobacteriaceae*. Emerg Infect Dis 2012; 18: 1503–7.

19. Yoon SH, Ha SM, Lim J et al. A large-scale evaluation of algorithms to calculate average nucleotide identity. Antonie Van Leeuwenhoek 2017; 110: 1281–6.

20. Meier-Kolthoff JP, Auch AF, Klenk HP et al. Genome sequence-based species delimitation with confidence intervals and improved distance functions. BMC Bioinformatics 2013; 14: 60.

21. Ciufo S, Kannan S, Sharma S et al. Using average nucleotide identity to improve taxonomic assignments in prokaryotic genomes at the NCBI. Int J Syst Evol Micr 2018; 68: 2386–92.

22. Miyoshi-Akiyama T, Hayakawa K, Ohmagari N et al. Multilocus sequence typing (MLST) for characterization of *Enterobacter cloacae*. PLoS One 2013; 8: e66358.

23. Gardner SN, Slezak T, Hall BG. kSNP3.0: SNP detection and phylogenetic analysis of genomes without genome alignment or reference genome. Bioinformatics 2015; 31: 2877–8.

24. Carattoli A, Zankari E, Garcia-Fernandez A et al. In silico detection and typing of plasmids using PlasmidFinder and plasmid multilocus sequence typing. Antimicrob Agents Chemother 2014; 58: 3895–903.

25. Robertson J, Nash JHE. MOB-suite: software tools for clustering, reconstruction and typing of plasmids from draft assemblies. Microb Genom 2018; 4: e000206.

26. Siguier P, Perochon J, Lestrade L et al. ISfinder: the reference centre for bacterial insertion sequences. Nucleic Acids Res 2006; 34: D32–6.

27. Aziz RK, Bartels D, Best AA et al. The RAST Server: rapid annotations using subsystems technology. BMC Genomics 2008; 9: 75.

28. Tanizawa Y, Fujisawa T, Nakamura Y. DFAST: a flexible prokaryotic genome annotation pipeline for faster genome publication. Bioinformatics 2018; 34: 1037–9.

29. Bortolaia V, Kaas RS, Ruppe E et al. ResFinder 4.0 for predictions of phenotypes from genotypes. J Antimicrob Chemother 2020; 75: 3491–500.

30. Moura A, Soares M, Pereira C et al. INTEGRALL: a database and search engine for integrons, integrases and gene cassettes. Bioinformatics 2009; 25: 1096–8.

31. Bertelli C, Laird MR, Williams KP et al. IslandViewer 4: expanded prediction of genomic islands for larger-scale datasets. Nucleic Acids Res 2017; 45: W30–W5.

32. Wick RR, Judd LM, Gorrie CL et al. Unicycler: Resolving bacterial genome assemblies from short and long sequencing reads. PLoS Comput Biol 2017; 13: e1005595.

33. Kolmogorov M, Yuan J, Lin Y et al. Assembly of long, error-prone reads using repeat graphs. Nat Biotechnol 2019; 37: 540–6.

34. Bush K, Jacoby GA. Updated functional classification of beta-lactamases. Antimicrob Agents Chemother 2010; 54: 969–76.

35. Naas T, Dortet L, Iorga BI. Structural and Functional Aspects of Class A Carbapenemases. Curr Drug Targets 2016; 17: 1006–28.

36. Uwamino Y, Kubota H, Sasaki T et al. Recovery of FRI-5 carbapenemase at a Japanese hospital where FRI-4 carbapenemase was discovered. J Antimicrob Chemother 2019; 74: 3390–2.

37. Adachi F, Sekizuka T, Yamato M et al. Characterization of FRI carbapenemase-producing *Enterobacter* spp. isolated from a hospital and the environment in Osaka, Japan. J Antimicrob Chemother 2021; 76: 3061–2.

38. Henriques I, Moura A, Alves A et al. Molecular characterization of a carbapenem-hydrolyzing class A beta-lactamase, SFC-1, from *Serratia fonticola* UTAD54. Antimicrob Agents Chemother 2004; 48: 2321–4.

39. Saavedra MJ, Peixe L, Sousa JC et al. Sfh-I, a subclass B2 metallo-beta-lactamase from a *Serratia fonticola* environmental isolate. Antimicrob Agents Chemother 2003; 47: 2330–3.

40. Henriques I, Juca Ramos RT, Barauna RA et al. Draft Genome Sequence of *Serratia fonticola* UTAD54, a Carbapenem-Resistant Strain Isolated from Drinking Water. Genome Announc 2013; 1: e00970–13.

41. Tijet N, Boyd D, Patel SN et al. Evaluation of the Carba NP test for rapid detection of carbapenemase-producing Enterobacteriaceae and *Pseudomonas aeruginosa*. Antimicrob Agents Chemother 2013; 57: 4578–80.

42. Boyd DA, Mataseje LF, Davidson R et al. *Enterobacter cloacae* Complex Isolates Harboring *bla*_NMC-A_ or *bla*_IMI-_Type Class A Carbapenemase Genes on Novel Chromosomal Integrative Elements and Plasmids. Antimicrob Agents Chemother 2017; 61: e02578–16.

43. Ho PL, Wang Y, Wing-Sze Tse C et al. Rapid Detection of Carbapenemase Production in Enterobacteriaceae by Use of a Modified Paper Strip Carba NP Method. J Clin Microbiol 2018; 56: e01110–17.

44. Zong Z, Feng Y, McNally A. Carbapenem and Colistin Resistance in *Enterobacter*: Determinants and Clones. Trends Microbiol 2021; 29: 473–6.

45. Wu W, Feng Y, Zong Z. Precise Species Identification for *Enterobacter*: a Genome Sequence-Based Study with Reporting of Two Novel Species, *Enterobacter quasiroggenkampii* sp. nov. and *Enterobacter quasimori* sp. nov. mSystems 2020; 5: e00527–20.

46. Chun J, Oren A, Ventosa A et al. Proposed minimal standards for the use of genome data for the taxonomy of prokaryotes. Int J Syst Evol Micr 2018; 68: 461–6.

47. Gomi R, Matsuda T, Yamamoto M et al. Characteristics of Carbapenemase-Producing Enterobacteriaceae in Wastewater Revealed by Genomic Analysis. Antimicrob Agents Chemother 2018; 62: e02501–17.

48. Brouwer MSM, Tehrani K, Rapallini M et al. Novel Carbapenemases FLC-1 and IMI-2 Encoded by an *Enterobacter cloacae* Complex Isolated from Food Products. Antimicrob Agents Chemother 2019; 63: e02338–18.

49. Meunier D, Findlay J, Doumith M et al. FRI-2 carbapenemase-producing *Enterobacter cloacae* complex in the UK. J Antimicrob Chemother 2017; 72: 2478–82.

50. Devi U, Khatri I, Kumar N et al. Draft Genome Sequence of Plant-Growth-Promoting Rhizobacterium *Serratia fonticola* Strain AU-AP2C, Isolated from the Pea Rhizosphere. Genome Announc 2013; 1: e01022–13.

51. Langille MGI, Hsiao WWL, Brinkman FSL. Detecting genomic islands using bioinformatics approaches. Nature Reviews Microbiology 2010; 8: 372–82.

52. Al-Hasani K, Rajakumar K, Bulach D et al. Genetic organization of the she pathogenicity island in *Shigella flexneri* 2a. Microb Pathog 2001; 30: 1–8.

53. Williams KP. Integration sites for genetic elements in prokaryotic tRNA and tmRNA genes: sublocation preference of integrase subfamilies. Nucleic Acids Res 2002; 30: 866–75.

54. Sullivan MJ, Petty NK, Beatson SA. Easyfig: a genome comparison visualizer. Bioinformatics 2011; 27: 1009–10.

55. Yu GC, Smith DK, Zhu HC et al. GGTREE: an R package for visualization and annotation of phylogenetic trees with their covariates and other associated data. Methods Ecol Evol 2017; 8: 28–36.

