## Supplementary Materials and methods, Supplementary Results, Supplemental Tables (S1, S5-S7), and Supplementary Figures (S1-S6) for "Emergence of rare carbapenemases (FRI, GES-5, IMI, SFC, and SFH-1) in Enterobacterales isolated from surface waters in Japan"

**Supplementary data**

**Supplementary Materials and methods**

**Antibiotic susceptibility testing**

Antibiotic susceptibility was assessed by microdilution using a Dry Plate Eiken (Eiken, Tokyo, Japan). The antibiotics used for the susceptibility testing included the following agents: ampicillin, piperacillin, ampicillin/sulbactam, amoxicillin/clavulanic acid, piperacillin/tazobactam, aztreonam, cefazolin, cefaclor, cefpodoxime, cefotaxime, ceftazidime, cefepime, cefoxitin, cefmetazole, flomoxef, doripenem, imipenem, meropenem, nalidixic acid, levofloxacin, ciprofloxacin, minocycline, tigecycline, gentamicin, tobramycin, amikacin, colistin, trimethoprim/sulfamethoxazole, and fosfomycin.

**Genome sequencing and assembly**

DNA was extracted from each isolate using a DNeasy Blood and Tissue kit (Qiagen, Hilden, Germany) for DNBSEQ short-read sequencing. Libraries were prepared with the MGIEasy FS PCR-free DNA library prep set (MGI Tech, Shenzhen, China), and paired-end sequencing (2×150-bp) was performed on DNBSEQ-G400RS. Reads were subsampled using seqtk (v1.3, https://github.com/lh3/seqtk), trimmed using fastp (v0.20.0),^1^ and assembled using Unicycler (v0.4.8) with the --no_correct option.^2^

Selected isolates (n=12) were also subjected to Oxford Nanopore Technologies (ONT) long-read sequencing. DNA was isolated as described previously,^3^ and libraries were prepared using the SQK-LSK109 kit (Oxford Nanopore Technologies, Oxford, UK) and the Short Read Eliminator XS kit (Circulomics, Inc., Baltimore, MD, USA). Sequencing was performed using ONT GridION X5 with a FLO-MIN106 flow cell. ONT reads were filtered using Filtlong (v0.2.0, https://github.com/rrwick/Filtlong) to remove reads with low quality or short length. *De novo* hybrid assemblies using trimmed DNBSEQ reads and filtered ONT reads were performed using Unicycler (v0.4.8)^2^ with the --no_correct option. Genomes that could not be completed by Unicycler were subjected to long-read assembly by Flye (v2.8)^4^ with the --plasmids option followed by long-read polishing using Medaka (v1.2.6, https://github.com/nanoporetech/medaka) and short-read polishing using Pilon (v1.23).^5^

**Supplementary Results**

**Comparison of *bla*_GES-5_-carrying plasmids**

A BLASTN search against the GenBank database using pJBIWA001_5 as a query sequence identified some closely related plasmids. pN260-3, which was carried by an *Enterobacter roggenkampii* strain detected in a patient with cholecystitis in Japan in 2019,^6^ showed the highest query coverage (97%) and also showed the high percent identity (>96%) (**Figure S1A**). pN260-3 also carried *bla*_GES-5_ but within a different class 1 integron structure (*intI1*-*bla*_GES-5_-aac(6*′*)-*31*-*intI1Δ*-*qacG2*-*aadA5*-*qacEΔ1*-*sul1*) (**Figure S1B**). The close relationship between plasmids pJBIWA001_5 and pN260-3 indicates potential interspecies dissemination of a plasmid carrying *bla*_GES-5_. Indeed, MOB-typer predicted that both pJBIWA001_5 and pN260-3 are mobilizable. Moreover, *bla*_GES-5_ was within a class 1 integron with intact IRi (inverted repeat at *intI1* end) and IRt (inverted repeat at *tni* end), which raises the possibility that *bla*_GES-5_ can be moved as a gene cassette or the structure from IRi to IRt can be moved by Tni proteins provided *in trans*.^7, 8^

**Supplementary Figures**

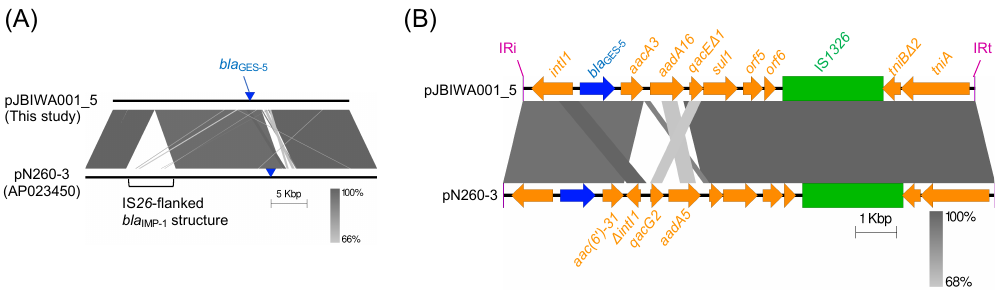

**Figure S1.** (A) Alignment of pJBIWA001_5 and pN260-3. Positions of *bla*_GES-5_ are indicated by blue triangles. The IS*26*-flanked *bla*_IMP-1_ structure (IS*26*-*tniR*-*bla*_IMP-1_-*tn*-*tn*-*aac(6')-I1*-*fosX*-*aac(6')-I1*-IS*26*)^6^ was only present in pN260-3. (B) Genetic contexts of *bla*_GES-5_ in pJBIWA001_5 and pN260-3. Blue arrows indicate *bla*_GES-5_, green boxes indicate insertion sequences, purple bars indicate IRi (inverted repeat at *intI1* end) and IRt (inverted repeat at *tni* end), and orange arrows indicate other coding sequences/genes.

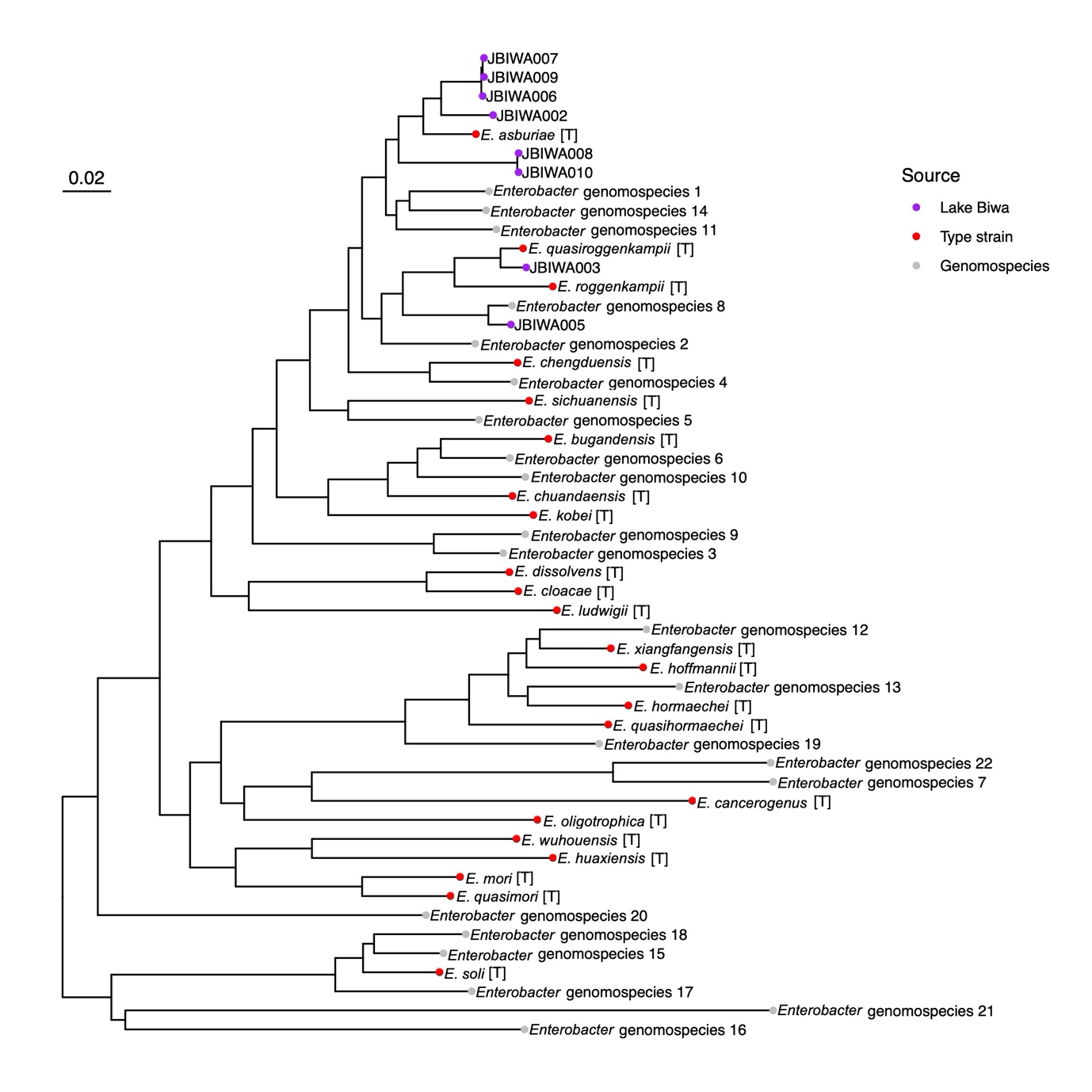

**Figure S2**. Phylogenetic tree of *Enterobacter* genomes. The tree was constructed based on 30,266 SNPs occurring in at least 80% of the genomes. The mid-point rooted tree was visualized with ggtree.^9^

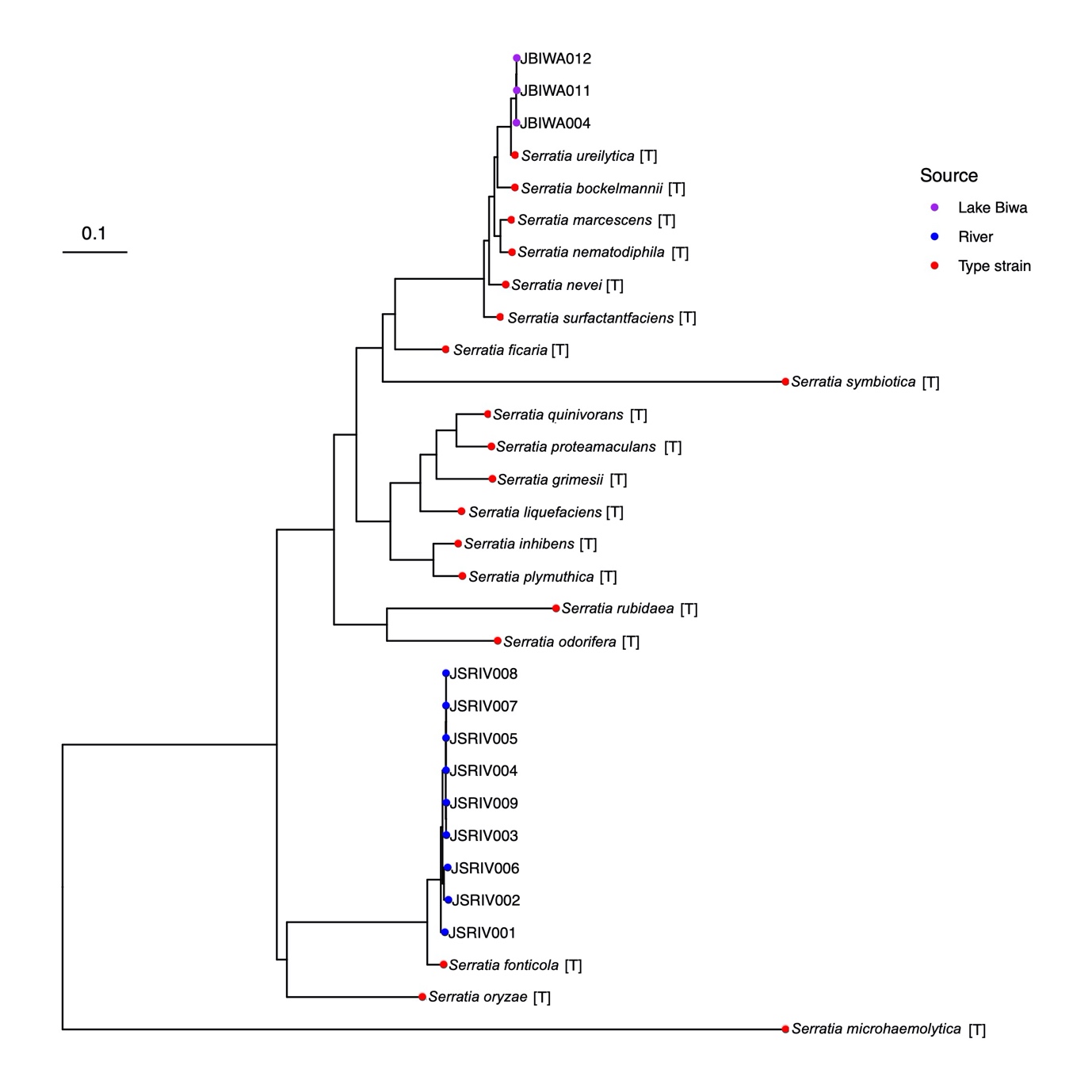

**Figure S3**. Mid-point rooted tree of *Serratia* genomes. The tree was constructed based on 9,430 SNPs occurring in at least 80% of the genomes.

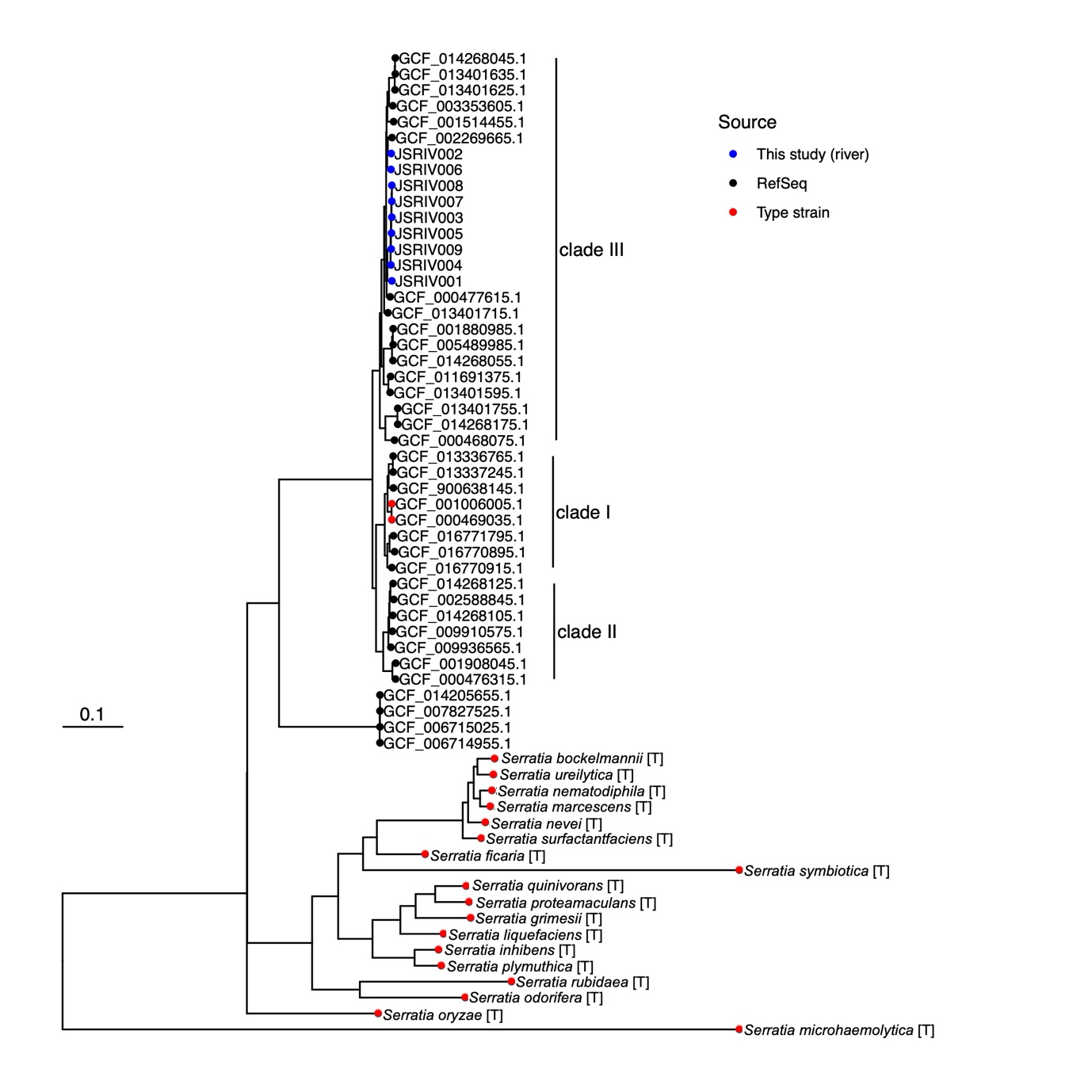

**Figure S4**. Mid-point rooted tree of RefSeq assemblies labelled *S. fonticola* and genomes of type strains of other *Serratia* species. The tree was constructed based on 16,369 SNPs occurring in at least 80% of the genomes. Four assemblies (GCF_006714955.1, GCF_006715025.1, GCF_007827525.1, GCF_014205655.1) were considered not to be *S. fonticola*.

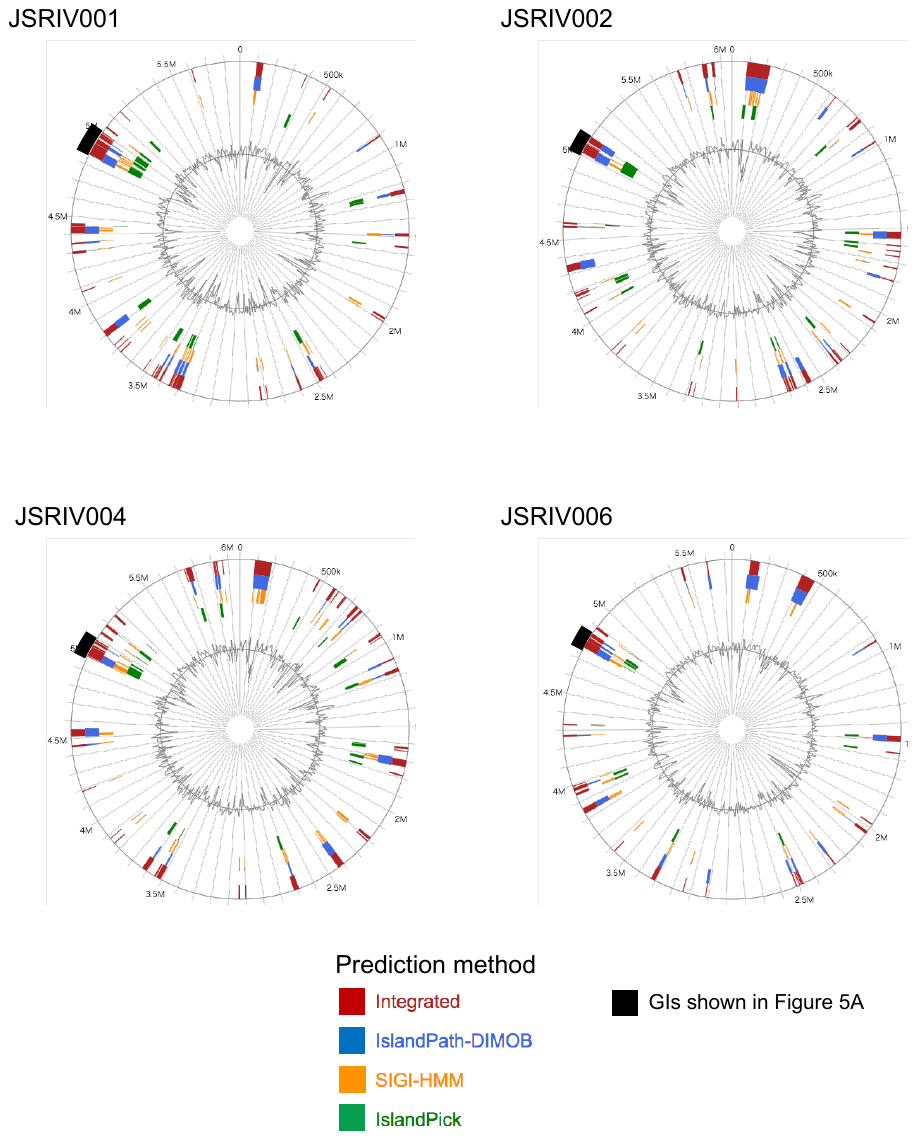

**Figure S5**. Genomic islands (GIs) predicted by IslandViewer 4. Four closed chromosomes and the predicted GIs are shown. IslandViewer 4 predicts GIs using three methods (IslandPath-DIMOB, SIGI-HMM and IslandPick) and also shows GIs that are predicted by one or more tools (called “Integrated” here). Putative GIs shown in **Figure 5A** are indicated by black bars.

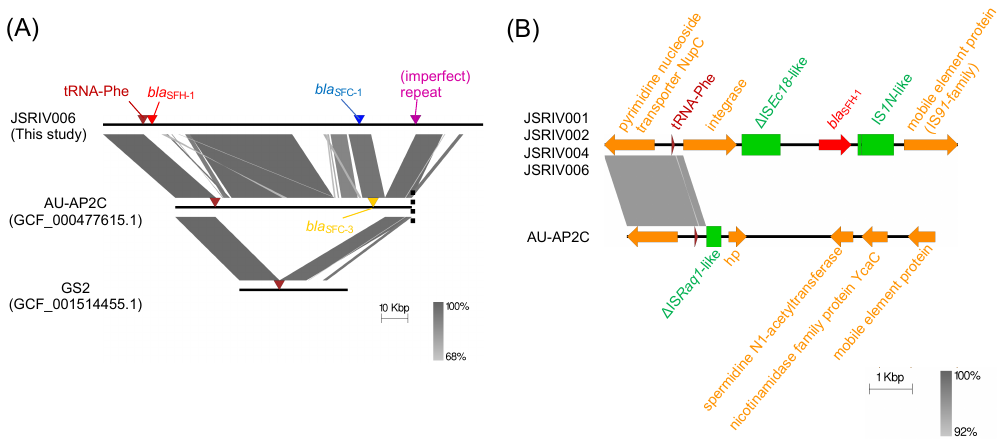

**Figure S6**. (A) Alignment of the *bla*_SFC-3_-carrying contig from the draft genome assembly of AU-AP2C against the chromosomes of JSRIV006 and GS2. JSRIV006 and GS2 are shown for the purpose of comparison. The vertical dashed line indicates a contig break. No (imperfect) repeat corresponding to the 3′-end of the tRNA-Phe gene was detected in AU-AP2C. (B) Comparison of genomic regions surrounding the tRNA-Phe genes.

**Supplementary Tables**

**Table S1**. Primers used for cloning experiments.^a^

| Target gene | Sequence (5′-3′) | Amplicon size (bp) |
| --- | --- | --- |
| *bla*_FRI-11_ | F-GCGATTTCTCATGGTATACTAGCC | 1,335 |
|  | R-TGCGCCATCGAATATACTGAT |  |
| *bla*_SFC-2_ | F-TGCAATTATGGAGAAAGGGCA | 1,232 |
|  | R-TGCGGCTTTAACTCAGCTGA |  |
| *bla*_IMI-22_ | F-ACATTTAATGGTAATCTGGCATGAA | 1,071 |
|  | R-CCATCACAATAAATTTATGGCTACAGA |  |
| *bla*_IMI-23_ | F-GCTACGTTCCTGAGGCTTCA | 1,099 |
|  | R-ACAGACGGTTAAAATGTGACGC |  |

^a^Primers were designed using Primer3 software^10^ to amplify the target carbapenemase gene and upstream region.

**Table S2**. Key features of Enterobacterales genomes sequenced to closure in this study (n = 11).

See the excel file.

**Table S3**. Key features of draft genomes of Enterobacterales isolates sequenced in this study (n = 10).

See the excel file.

**Table S4**. Resistance phenotypes of Enterobacterales isolates.

See the excel file.

**Table S5**. MICs (mg/L) of different antibiotics for the isolates analysed in this study.

| Antibiotic | *E. coli* DH5α  (pCR2.1-TOPO*-bla*_FRI-11_) | *E. coli* DH5α  (pCR2.1-TOPO*-bla*_SFC-2_) | *E. coli* DH5α  (pCR2.1-TOPO-*bla*_IMI-22_) | *E. coli* DH5α  (pCR2.1-TOPO-*bla*_IMI-23_) | *E. coli* DH5α  (pCR2.1-TOPO)^a^ | *E. coli* DH5α |
| --- | --- | --- | --- | --- | --- | --- |
| Ampicillin | >16 | >16 | >16 | >16 | >16 | ≤8 |
| Piperacillin | >16 | >16 | >16 | >16 | >16 | ≤16 |
| Aztreonam | ≤4 | >8 | ≤4 | >8 | ≤4 | ≤4 |
| Cefazolin | >16 | >16 | 16 | >16 | 16 | ≤2 |
| Cefpodoxime | ≤1 | >4 | ≤1 | 2 | ≤1 | ≤1 |
| Cefepime | ≤2 | 4 | ≤2 | ≤2 | ≤2 | ≤2 |
| Cefotaxime | ≤1 | ≤1 | ≤1 | ≤1 | ≤1 | ≤1 |
| Ceftazidime | ≤1 | 4 | ≤1 | ≤1 | ≤1 | ≤1 |
| Cefmetazole | 2 | 8 | 2 | 2 | 0.5 | 1 |
| Doripenem | ≤1 | >2 | ≤1 | ≤1 | ≤1 | ≤1 |
| Meropenem | ≤0.12 | >8 | ≤0.12 | 0.5 | ≤0.12 | ≤0.12 |
| Imipenem | ≤1 | >8 | ≤1 | >8 | ≤1 | ≤1 |

^a^pCR2.1-TOPO contains an ampicillin resistance gene (*bla*_TEM-116_) and a kanamycin resistance gene (*aph(3')-IIa*).

**Table S6**. Pairwise average nucleotide identity (ANI) values within and between *S. fonticola* clades.^a^

|  | Clade I | Clade II | Clade III |
| --- | --- | --- | --- |
| Clade I | 98.25±0.58  (97.69-99.98) | 95.51±0.18  (95.13-95.82) | 95.58±0.15  (95.17-95.83) |
| Clade II |  | 98.01±1.04  (96.85-99.15) | 95.25±0.19  (94.73-95.54) |
| Clade III |  |  | 98.08±0.83  (96.65-100.00) |

^a^Mean and standard deviation are shown above and the minimum and maximum values are shown in parentheses.

**Table S7**. Pairwise digital DNA-DNA hybridization (dDDH) values for the type and reference strains vs genomes from each clade.^a^

|  | Clade I | Clade II | Clade III |
| --- | --- | --- | --- |
| GCF_001006005.1  (Clade I, type strain of *S. fonticola*) | 86.83±6.33  (81.70-99.80) | 64.29±1.09  (62.60-65.40) | 65.00±0.90  (62.40-66.20) |
| GCF_002588845.1  (Clade II, reference strain^b^) | 65.64±0.30  (65.30-66.10) | 86.65±9.07  (74.80-93.00) | 63.45±0.74  (61.60-64.10) |
| GCF_001514455.1  (Clade III, reference strain^b^) | 65.18±0.43  (64.50-65.60) | 62.49±0.90  (61.1-63.5) | 83.27±4.12  (74.20-87.10) |

^a^Mean and standard deviation are shown above and the minimum and maximum values are shown in parentheses.

^b^Reference strains were selected based on completeness of the genomes.

**References**

1.	Chen, S.; Zhou, Y.; Chen, Y.; Gu, J., fastp: an ultra-fast all-in-one FASTQ preprocessor. *Bioinformatics* **2018,** *34* (17), i884-i890.

2.	Wick, R. R.; Judd, L. M.; Gorrie, C. L.; Holt, K. E., Unicycler: Resolving bacterial genome assemblies from short and long sequencing reads. *PLoS Comput Biol* **2017,** *13* (6), e1005595.

3.	Morita, H.; Kuwahara, T.; Ohshima, K.; Sasamoto, H.; Itoh, K.; Hattori, M.; Hayashi, T.; Takami, H., An improved DNA isolation method for metagenomic analysis of the microbial flora of the human intestine. *Microbes Environ* **2007,** *22* (3), 214-222.

4.	Kolmogorov, M.; Yuan, J.; Lin, Y.; Pevzner, P. A., Assembly of long, error-prone reads using repeat graphs. *Nat Biotechnol* **2019,** *37* (5), 540-546.

5.	Walker, B. J.; Abeel, T.; Shea, T.; Priest, M.; Abouelliel, A.; Sakthikumar, S.; Cuomo, C. A.; Zeng, Q.; Wortman, J.; Young, S. K.; Earl, A. M., Pilon: an integrated tool for comprehensive microbial variant detection and genome assembly improvement. *PLoS One* **2014,** *9* (11), e112963.

6.	Umeda, K.; Nakamura, H.; Fukuda, A.; Matsumoto, Y.; Motooka, D.; Nakamura, S.; Yasui, Y.; Yoshida, H.; Kawahara, R., Genomic characterization of clinical *Enterobacter roggenkampii* co-harbouring *bla*_IMP-1_- and *bla*_GES-5_-encoding IncP6 and *mcr-9*-encoding IncHI2 plasmids isolated in Japan. *J Glob Antimicrob Resist* **2020,** *24*, 220-227.

7.	Partridge, S. R., Analysis of antibiotic resistance regions in Gram-negative bacteria. *FEMS Microbiol Rev* **2011,** *35* (5), 820-55.

8.	Partridge, S. R.; Kwong, S. M.; Firth, N.; Jensen, S. O., Mobile Genetic Elements Associated with Antimicrobial Resistance. *Clin Microbiol Rev* **2018,** *31* (4), e00088-17.

9.	Yu, G. C.; Smith, D. K.; Zhu, H. C.; Guan, Y.; Lam, T. T. Y., GGTREE: an R package for visualization and annotation of phylogenetic trees with their covariates and other associated data. *Methods Ecol Evol* **2017,** *8* (1), 28-36.

10.	Untergasser, A.; Cutcutache, I.; Koressaar, T.; Ye, J.; Faircloth, B. C.; Remm, M.; Rozen, S. G., Primer3--new capabilities and interfaces. *Nucleic Acids Res* **2012,** *40* (15), e115.
